# Investigating the Effects of *APOE* Genotype on Intracellular Cholesterol and the Endolysosomal System in the Aging Mouse Brain

**DOI:** 10.1101/2025.11.11.687891

**Authors:** Sherida M. de Leeuw, Tal Nuriel

## Abstract

Individuals who possess the ε4 allele of apolipoprotein E (*APOE*) are at a significantly increased risk for developing Alzheimer’s disease (AD). However, the precise reason for this association is not fully understood. Beyond its effects on amyloid and tau, *APOE* also influences fundamental cellular processes in the brain, including cholesterol trafficking between cells and within the endolysosomal system, which may be a critical component of the *APOE4*-associated vulnerability to AD. Here, we examined how *APOE* genotype, sex, and aging alter intracellular cholesterol processing and the endolysosomal system in the mouse brain. Using the novel cholesterol-binding probe D4H*-mCherry, we quantified intracellular cholesterol levels, the levels of early endosomes (Rab5), late endosomes (CD63), and lysosomes (LAMP1), and the colocalization of cholesterol with these endolysosomal compartments. This analysis was performed in the cortex, hippocampus, and entorhinal cortex of young, middle-aged, and old *APOE2*, *APOE3*, and *APOE4* mice. Our analysis revealed region-specific changes in intracellular cholesterol and the endolysosomal system in response to aging, sex, and *APOE* genotype. Notably, young *APOE4* mice showed reduced cholesterol within early and late endosomes, but increased lysosomal abundance, suggesting impaired cholesterol processing. These APOE4-specific effects were less apparent in older animals. These effects were strongly modified by sex, with female *APOE4* mice exhibiting elevated lysosomal cholesterol in the hippocampus and entorhinal cortex at old age, indicating sex-dependent susceptibility. Together, these results reveal that *APOE* genotype, age, and sex interact to influence endolysosomal cholesterol homeostasis, with female *APOE4* mice showing the greatest dysregulation. These findings suggest that early and region-specific endolysosomal defects may contribute to the heightened AD risk associated with *APOE4*, particularly in females.

## INTRODUCTION

Possession of the ε4 allele of apolipoprotein E (*APOE*) is the primary genetic risk factor for the late-onset, sporadic form of Alzheimer’s disease (AD), while possession of the ε2 allele confers resistance to AD. Although extensive research has been performed on *APOE* and AD, the precise cause of the increased risk of AD in *APOE4* carriers remains unclear. Importantly, while most research has focused on *APOE4*’s ability to increase AD pathology in the brain, this process likely begins decades before any AD pathology is present, with *APOE4* expression subtly altering key physiological processes in the brain, the effect of which accumulates with age.

During normal physiology, the APOE protein plays a vital role in the transport of cholesterol and other lipids through the bloodstream, as well as within the brain (Han, 2004; Holtzman et al., n.d.; Mahley & Rall, 2000). In the brain, APOE is primarily produced by astrocytes, where lipidation of APOE occurs extracellularly with the aid of ABCA7, followed by delivery of the lipidated APOE to neurons and other cell-types. However, the APOE4 variant has been shown to undergo poorer lipidation compared than APOE2 and APOE3 (Heinsinger et al., 2016; Hu et al., 2015), resulting in less lipid cargo being delivered to neurons. Importantly, neurons endocytose the lipidated APOE via binding to one of several lipoprotein receptors, with APOE2 having less binding affinity, particularly for the low density lipoprotein (LDL) and LDL receptor-related protein 1 (LRP1) receptors than APOE3 and APOE4 (Kowal et al., 1990; Schneider et al., 1981). Within the endolysosomal system, the lipids are cleaved from APOE and distribution within the neurons for utilization in a variety of cellular functions.

Previous studies, including our own, have shown that the APOE4 variant can increase the amount of intracellular cholesterol inside cells (de Leeuw et al., 2022; TCW et al., 2022), which is most likely due to aberrant endolysosomal trafficking of APOE4 (DeKroon & Armati, 2001; Gong et al., 2002; Heeren et al., 2004; Nuriel et al., 2017). We have also observed an increase in early endosomal and lysosomal compartments in aged *APOE4* mice, compared to *APOE3 (Nuriel et al., 2017)*. Interestingly cholesterol metabolism in the brain changes with age. Studies have shown that cholesterol levels in the brain decrease with age, especially in the hippocampus, and neuronal membranes (Nunes et al., 2022; Yu et al., 2020). This is paired with both a decrease in cholesterol synthesis, and increased levels of 24-hydroxylase – the enzyme that facilitates cholesterol removal from the brain. Human studies have shown that a critical time window for changes in cholesterol metabolism is around 55 years of age. Importantly, cholesterol is also a building block for hormones such as estrogen (Cui et al., 2013). This coincides with menopause in women, and the significant loss of estrogen production, which can also be paired with cognitive symptoms. Women are at higher risk for AD, and the *APOE4* allele is a higher risk factor for women compared to men (Belloy et al., 2023). Altogether, changes in cholesterol metabolism in combination with APOE4 function have been suggested to play a fundamental role in AD pathobiology (Maioli et al., 2025). However, there is little understanding of the patterns and mechanisms of endolysosomal cholesterol changes throughout aging, dependent on *APOE* genotype and sex.

Previous studies showing aberrant endolysosomal cholesterol trafficking by APOE4 have primarily been performed in cultured cells. This is partly due to the difficulty of visualizing and measuring intracellular cholesterol in tissues. Filipin III, the primary research tool used to measure cellular cholesterol, binds strongly to plasma membrane cholesterol and is also prone to photobleaching during imaging, making it difficult to use for intracellular cholesterol investigations in tissues. In order to overcome this obstacle, we optimized an intracellular cholesterol staining protocol (de Leeuw & Nuriel, 2024), using the novel cholesterol-binding reagent D4H*-mCherry. D4 is the fourth domain (the cholesterol binding domain) of the Perfingolysin O (PFO) toxin, and it has previously been adapted (D434S, Y415A, A463W) to generate a higher cholesterol affinity variant called D4H* (Liu et al., 2016; Maekawa & Fairn, 2015). Furthermore, tissue fixation prevents plasma membrane binding of PFO-derived reagents (Reid et al., 2004), thus allowing us to use D4H*-mCherry for specific investigation of intracellular cholesterol in the mouse brain.

In this study, we performed a comprehensive analysis of intracellular cholesterol trafficking in the mouse brain, with a specific focus on the impact of *APOE* genotype, sex, and aging on intracellular cholesterol levels and cholesterol localization within different endo-lysosomal compartments. To achieve this, we stained fixed brain tissues from young, middle-aged, and old *APOE2*, *APOE3*, and *APOE4* mice with D4H*-mCherry and with antibodies for one of three endolysosomal markers: Rab5 (early endosomes), CD63 (late endosomes), or LAMP1 (lysosomes). We then imaged the stained tissues using high throughput confocal microscopy, followed by detailed analyses using Imaris. These analyses revealed a complicated, but illuminating picture of how differential *APOE* isoform expression alters intracellular cholesterol trafficking, with important mediation of sex and age.

## METHODS

### APOE mice

The human *APOE* mice used in this study were developed by the Cure Alzheimer’s Fund at Taconic Biosciences (Huynh, 2019). All mice used in this study were treated in accordance with the National Institutes of Health Guide for the Care and Use of Laboratory Animals and approved by the McLaughlin Research Institute’s Institutional Animal Care and Use Committee (IACUC). For this study, we utilized 5-6 month-old (young), 15-16 month-old (middle-aged), and 24-26 month-old (old) male and female *APOE2/2*, *APOE3/3*, and *APOE4/4* mice, 5 mice per sex per genotype for each age group. Mice were euthanized via transcardial perfusion with 10% formalin and ice-cold PBS, and then the brains were drop-fixed for an additional 24 hours in 10% formalin, before transferring to 30% sucrose.

### D4H*-mCherry treatment and endolysosomal immunocytochemistry

The D4H*-mCherry treatments and endolysosomal stainings were performed as described previously (de Leeuw & Nuriel, 2024). The stainings were performed over the course of three separate experiments, one for each of the three endolysosomal markers: Rab5 (early endosomes), CD63 (late endosomes), or LAMP1 (lysosomes). In addition, neuronal soma were stained with NeuN and nuclei were stained with DAPI (25µg/mL, ref: 62248, Invitrogen). Antibody details are listed below.

### Antibody table

| Target | Host animal | Reference number | Company | Dilution |
| --- | --- | --- | --- | --- |
| NeuN | Guinea pig | 266004 | Synaptic Systems | 1:500 |
| mCherry | Rabbit | GTX128508 | GeneTex | 1:500 |
| Rab5 | Rabbit | ab18211 | Abcam | 1:1000 |
| CD63 | Rabbit | ab217345 | Abcam | 1:500 |
| Lamp1 | Rat | 121602 | Biolegend | 1:500 |
| Biotin anti-Rabbit | Goat | 65-6140 | Invitrogen | 1:400 |
| AlexaFluor555 anti-Guinea pig | Goat | A21435 | Invitrogen | 1:1000 |
| AlexaFluor647 anti-Rabbit | Goat | A21245 | Invitrogen | 1:1000 |
| AlexaFluor488 anti-Rabbit | Goat | A11034 | Invitrogen | 1:1000 |
| AlexaFluor488 anti-Rat | Goat | A11006 | Invitrogen | 1:1000 |
| Fluorescein Tyramide |  | 6456 | Tocris | 7.5µM |

### Confocal imaging

Three brain regions per section were imaged: entorhinal cortex (EC), CA1 region of the hippocampus (HIPP), and neocortex (CTX). Five images per region were captured, totaling 15 images per section. Images were captured using a CSU-W1-Yokogawa spinning disk confocal microscope (Nikon), pinhole radius 25, with a high-sensitivity Prime BSI sCMOS camera and Perfect Focus System 4 to prevent focus drift. Binning: 1x1. Bit depth: 16-bit. Dimensions: 51 x Z-steps at 0.2μm, resulting in a 10μm Z-stack, and four fluorescent channels: AF647, AF568, GFP, and DAPI. The objective used was a 100x/1.45 NA oil Plan-Apochromat λ (Zeiss).

### Imaris image analysis and data export

Raw .dn2 file images were imported into Imaris, with voxel size 0.065x0.065x0.200, and batch processes for each staining experiment. DAPI nuclei were identified using the objects module and segmented using the machine learning option. Similarly, NeuN-AF555 labeled neuronal soma were identified using the object module and segmented with the machine learning option. D4H*-AF647 spots were identified using the spots module, with an estimated XY diameter of 0.5µm, estimated Z diameter of 1 µm, and background subtraction was applied. To select the spots that were regarded as true spots a Quality spot filter of above 25 was applied. Rab5- and CD63-AF488 labeled spots were defined using the spots module, with an estimated XY diameter of 0.6µm or 0.65µm, estimated Z diameter of 1.2µm or 1.3µm, respectively, and background subtraction was applied. For Rab5-AF488 spots, the Quality filter was set at above 140, and for CD63-AF488 at above 50. Colocalized spots were defined by Rab5- or CD63-AF488 spots located at a distance of 0.2µm or below from D4H*-AF647 spots. Lysosomal Lamp1-AF488 had a more amorphous patterns and therefore was identified using the objects module instead of spots. Segmentation was run based on fluorescent intensity, smoothing enabled, background subtraction applied, and the Diameter Of Largest Sphere set to 0.9µm. Lamp1-AF488 objects were labeled as colocalized with D4H*-AF647 spots if the Shortest Distance to Objects was 0.0µm or below. Lastly, to identify spots or lysosomal objects within the neuronal object volume, a cut off distance to neuron object threshold of 0.0µm or below was applied. After batch processing, the measurements were exported to .CSV files.

### Data analysis with R

For each staining experiment, at each age group, the following metrics were calculated per image and graphed: 1. Total number of Rab5 spots | CD63 spots | Lamp1 objects/Number of DAPI nuclei, 2. (Total number of D4H* spots colocalized with Rab5 spots | CD63 spots | Lamp1 objects/Total number of D4H* spots)/Number of DAPI nuclei, 3. Neuronal number of Rab5 spots | CD63 spots | Lamp1 objects/Neuronal volume, 4. (Neuronal number of D4H* spots colocalized with Rab5 spots | CD63 spots | Lamp1 objects/Neuronal number of D4H* spots)/Neuronal volume. These metrics were graphed per sex per age group per genotype (N=75 images), or graphed per brain region specifically (N=15 images). To quantify D4H* spots, the data from three experiments per age group were taken together and graphed: 1. Total number of D4H* spots/Number of DAPI nuclei, and 2. Neuronal number of D4H* spots/Neuronal volume. This resulted in 3 repeated measures of N=75 images per condition in total or graphed per brain region specifically (N=15 images).

### Statistical analysis

Statistical analyses were performed to test the effect of *APOE* genotype (ε2/ε2, ε3/ε3, or ε4/ε4), *Age* (5-6 months (Young), 15-17 months (Middle age), or 24-26 months (Old), and *sex* (male or female). To preprocess the data before statistical analysis, data sets from each experiment were first normalized by z-scoring. Outliers were pruned based on Interquartile range-based outlier analysis, applying the Tukey 1.5×IQR rule. For each metric, the data were graphed as box plots grouped by *age*, sex, *APOE* genotype, or combinations of variables were plotted based on pooled data. Linear mixed-effects model: DV ∼ Sex × APOE × Age + (1|ID). Factors were coded with Young (Age), Male (Sex), and the *APOE* reference level as baselines. Wald χ² tests (HC3-robust) evaluated sex main effects, age main effect, *APOE* main effect, simple *age* effects within *sex*, Sex×Age interaction, *APOE*×Age interaction, Sex×*APOE* interaction, and three-way Sex×*APOE*×Age interaction. P-values were FDR-corrected across contrasts. Significant p-values (α = 0.05) were displayed on the graphs with significance bars, or interaction curves superimposed on the boxplots. For tests within one age group, the model was adapted. Linear mixed-effects model: DV ∼ Sex × APOE + (1|ID); Male and *APOE2* are reference levels. Wald χ² tests (HC3-robust) assessed *APOE* contrasts within Sex, and Sex differences within each *APOE* level. P-values were FDR-corrected across tests.

## RESULTS

### Overall effect on cholesterol levels

As an initial analysis of the data, we measured the effects of aging, *APOE* genotype, and sex on the overall cholesterol levels in our samples. For this analysis, data from the D4H*-mCherry positive spots were pooled from 3 experiments per age group, introducing a significant amount of technical variability. However, this method of data analysis indirectly tests the robustness of this method in identifying differences in cholesterol in brain sections.

Assessing the overall effects of aging, *APOE* genotype, and sex on total and neuron-specific D4H*-mCherry spots revealed no main effects exerted by these variables. Pairwise contrasts between sexes (Male vs Female, adjusted for *APOE* and *Age*) showed that there was a significant increase in total and neuronal cholesterol in female CTX compared to male CTX (**Fig. 1A-B**). This effect was region specific for the cortex, showing that cholesterol in females is higher than in males. There were no further pairwise differences in cholesterol based on *Age* or *APOE* status (**Fig. S1**).

**Figure 1.**
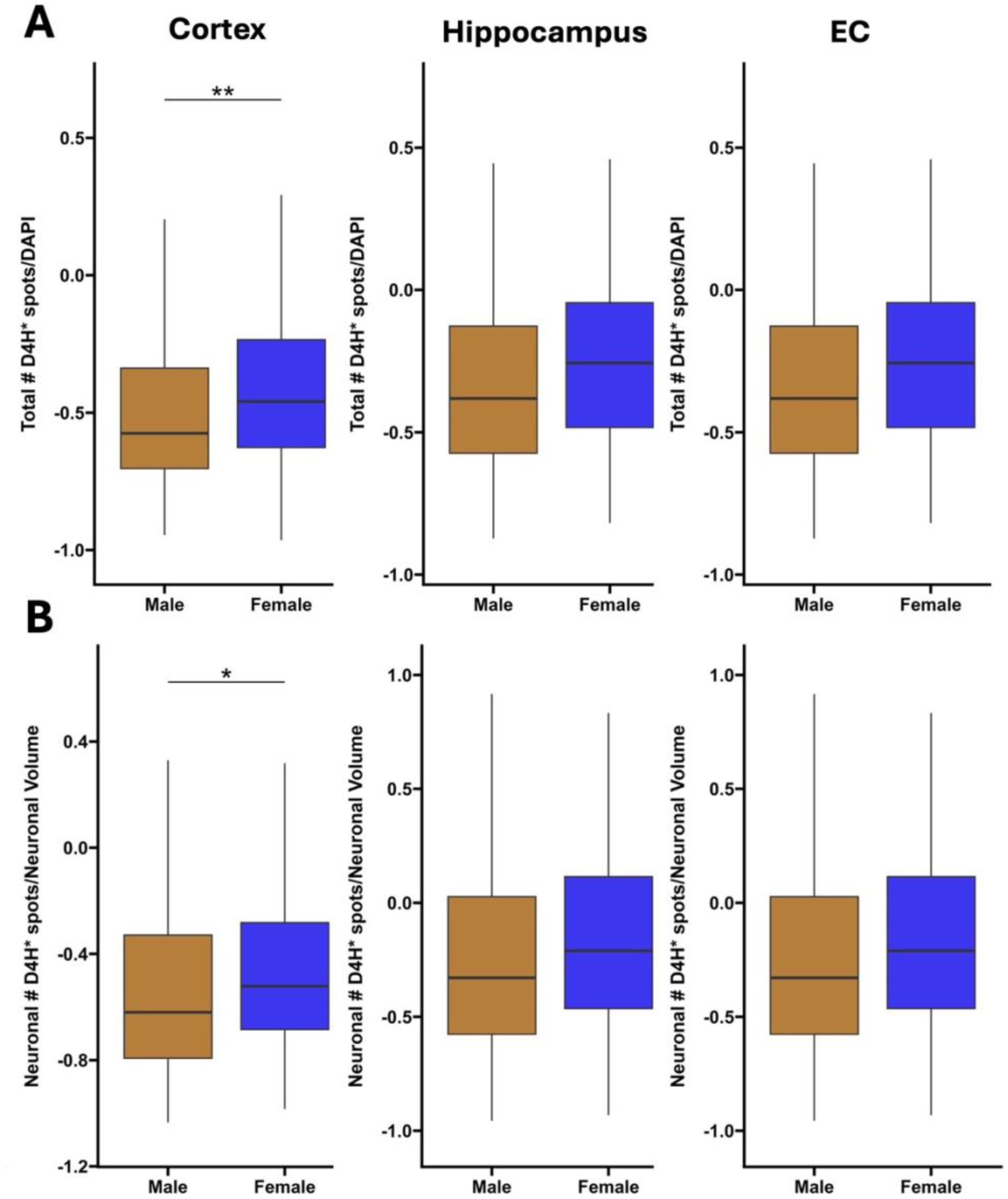
Cholesterol levels in males and females. **A)** Total levels of cholesterol, represented by box plots of total number of D4H* spots, normalized to DAPI nuclei. Comparisons per brain regions, cortex, hippocampus, EC. Male (ochre fill) and female (blue fill). **B)** Neuronal levels of cholesterol box plots, representing neuronal D4H* spots normalized for the NeuN neuronal volume. Statistics: data from individual experiments were Z-scored before pooling. Linear mixed-effects model was fitted for each variable using *sex*, *age*, and *APOE* genotype as fixed effects. Estimated marginal means for *sex*, and pairwise comparison between sexes, p<0.05: *, p<0.01:**. N = 5 animals per *sex*, *age*, *APOE* genotype. *Number of independent experiments: 3*.

### Effect of aging on the endolysosomal system and endolysosomal cholesterol

We next asked whether there were any overall aging, sex, or *APOE* effects in the endolysosomal or endolysosomal-cholesterol analyses from each region. These results are displayed in **Table 1**. All three main effects were generally small, but effects on both endolysosomal numbers and endolysosomal-cholesterol colocalization were observed. Levels of Rab5-containing early endosomes (EEs) were not affected by *age* (**Fig. 2A**). We observed a small *age* effect on the total number of CD63-containing late endosomes (LEs) in the CTX (**Fig. 2B**). This was driven by an increase of CD63 spots from the old age group. For LAMP1-containing lysosomes (LY), we observed a small main effect of *age* exclusive in the EC region (**Fig. 2C**). This suggests that aging alone has some effect on LE’s and LY’s, but not EE’s.

**Figure 2.**
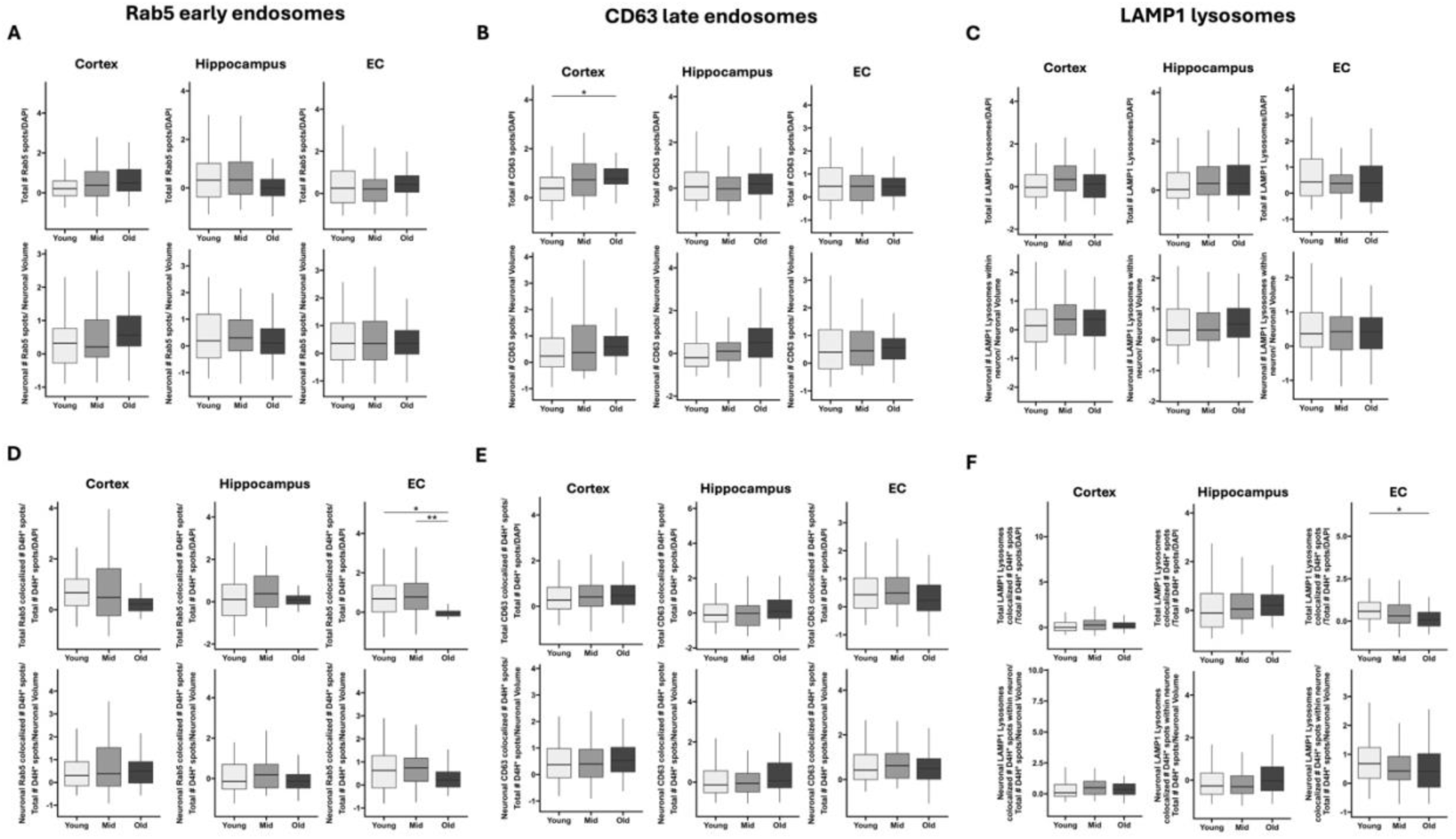
The main effect of *age* on early endosomes (EE), late endosomes (LE), lysosomes (LY), and cholesterol associated with EE’s, LE’s, and LY’s. **A**) Box plots representing total number of Rab5 spots, normalized to number of DAPI nuclei. *Lower row*: box plots representing neuronal number of Rab5 spots normalized to the total NeuN neuronal volume. The same metrics but different markers are depicted in plots **B)**, and **C)**. **B)** CD63 spots, and **C)** Lamp1 particles. **D)** Box plots representing total Rab5 colocalized D4H* spots, normalized to the total number of D4H* spots and subsequently normalized to the number of DAPI nuclei. *Lower row*: box plots representing neuronal number of Rab5 colocalized D4H* spots, to normalized to the total number of D4H* spots, and subsequently to the total NeuN neuronal volume. The same metrics but different markers are depicted in plots **D)**, and **F)**. **E)** CD63 spots, and **F)** Lamp1 particles. The data is grouped by age; legend in the figure, and by APOE genotype; *APOE2*, *APOE3*, *APOE4*. *Statistics:* data from individual experiments were Z-scored before pooling. Linear mixed-effects model was fitted for each variable using *sex*, *age*, and *APOE* genotype as fixed effects. Estimated marginal means for *age*, and pairwise comparison between age groups, p<0.05: *, p<0.01:**. N = 5 animals per sex, age, *APOE* genotype.

**Table 1.**
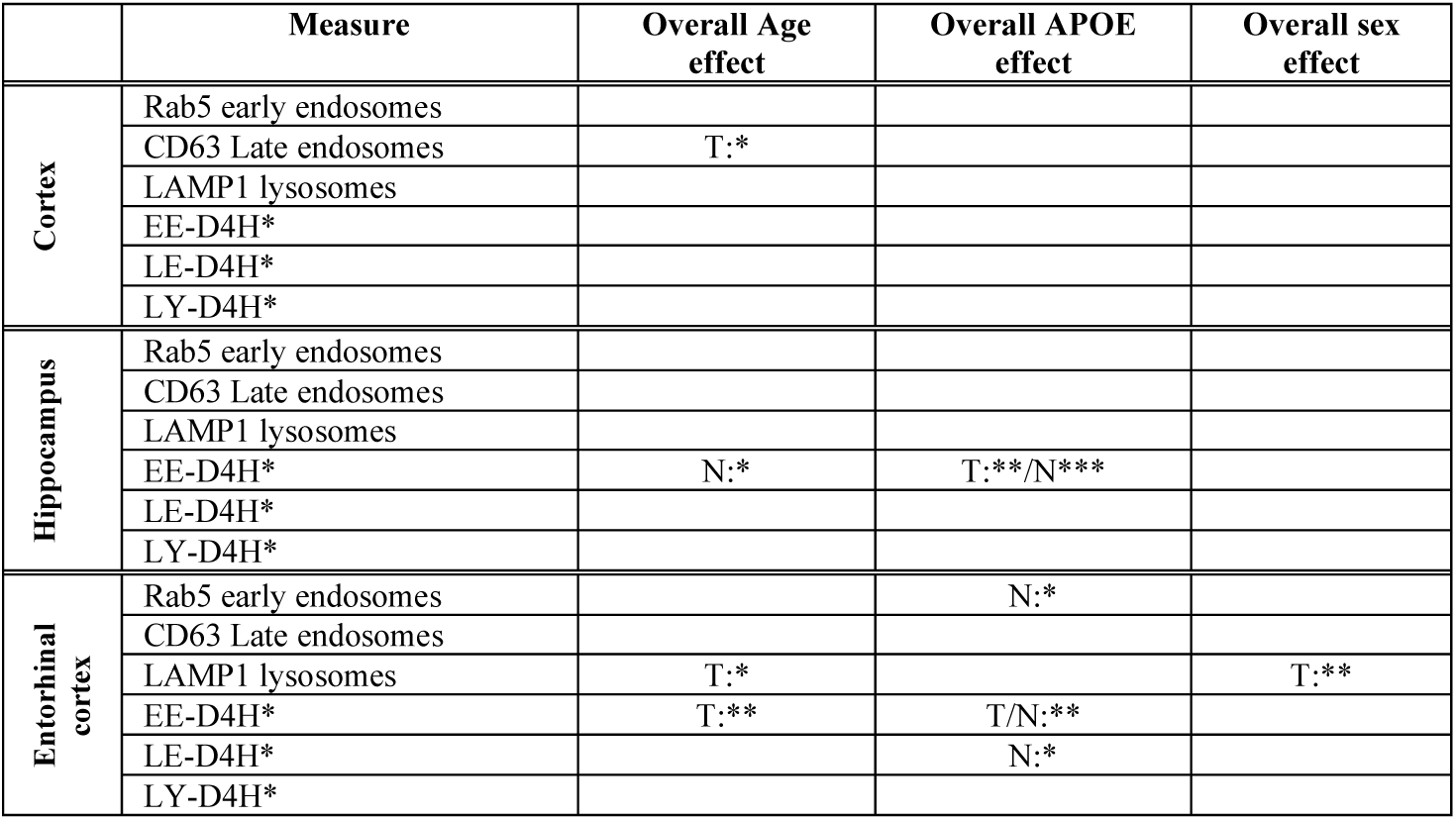
Overall effects of aging, *APOE* genotype, and sex on the endolysosomal system and endolysosomal cholesterol.

In terms of cholesterol colocalization, we observed an overall effect of *age* on total early endosomal cholesterol (EE-Ch), and neuronal EE-Ch (N-EE-Ch) in the EC and HIPP respectively, driven primarily by a decrease in T-EE-Ch in the old age group in the EC (**Fig. 2D**). This indicates that cells other than neurons play a role in the decrease of EE-Ch in the EC specifically. There was no overall *age* effect observed for LE-cholesterol (LE-Ch) in any of the brain regions (**Fig. 2E**). Lastly, for lysosomal cholesterol (LY-Ch), we did not observe a main effect of age, even though pairwise analyses (corrected for *APOE* and sex) did show a small decrease with age in T-LY-Ch in the EC exclusively (**Fig. 2F**). These data indicate that age is not a main driver of cholesterol accumulation in the endolysosomal system, but on the other hand does account for some small decreases with age specifically in the EC.

### Effect of *APOE* genotype and aging on the endolysosomal system and endolysosomal cholesterol

The primary goal of our study was to investigate the role of *APOE* genotype on endolysosomal cholesterol in the brain. In general, the main effect of *APOE* genotype on endolysosomes and endolysosomal cholesterol (controlled for by age and sex) were stronger than the aging effects that were shown in **Table 1**. In terms of an overall *APOE genotype* effect, none of the EE’s, LE’s, LY’s, or associated cholesterol, were affected in the cortex. Total and neuronal EE-Ch (T-EE-Ch and N-EE-Ch) both in the HIPP and EC were strongly affected by *APOE* genotype, corroborated by pairwise testing showing a decrease in T/N-EE-Ch in the EC with *APOE* genotype (*APOE2*>*E3*; *APOE2*>*E4*) (**Supplementary Fig. S2D**). Furthermore, an *APOE* effect was observed on N-LE-Ch levels in the EC (**Supplementary Fig. S2B**).

In addition to the main effect, we measured the *APOE genotype* x *age* interaction effects (**Table 2**). For EE’s, we observed a significant interaction effect between *APOE genotype* x *age* exclusively on N-EE levels in the EC, which was driven by neuronal EEs changing with age differently in *APOE4* vs. *APOE2* and *APOE4* vs. *APOE3* brains, but no difference between *APOE3* and *APOE2* (**Fig. 3A**). There were no significant interaction effects between *APOE x age* in LE levels (**Fig. 3B**). On the other hand, N-LY levels in the hippocampus showed a strong interaction between *APOE x age*, on all levels: *APOE3* vs *APOE2*; *APOE4* vs *APOE2*; *APOE4* vs *APOE3* (**Fig. 3C**). This was driven by Y<O in *APOE2*, Y<M>O in *APOE3*, and Y>M in *APOE4*, as well as *E2*<*E3*; *E2*<*E4* at young age, *E2*<*E3*; *E3*>*E4* at middle age, and *E2*>*E3* at old age. These data indicate that while neither *APOE* genotype nor age had any main effects on hippocampal N-LY levels, the interaction between age and *APOE* strongly affects lysosomal levels specifically in hippocampal neurons. Total LY levels in the EC also showed a small *APOE genotype* x *age* interaction effect, dependent on the difference between *APOE2* vs *APOE3* (**Fig. 3C**).

**Figure 3.**
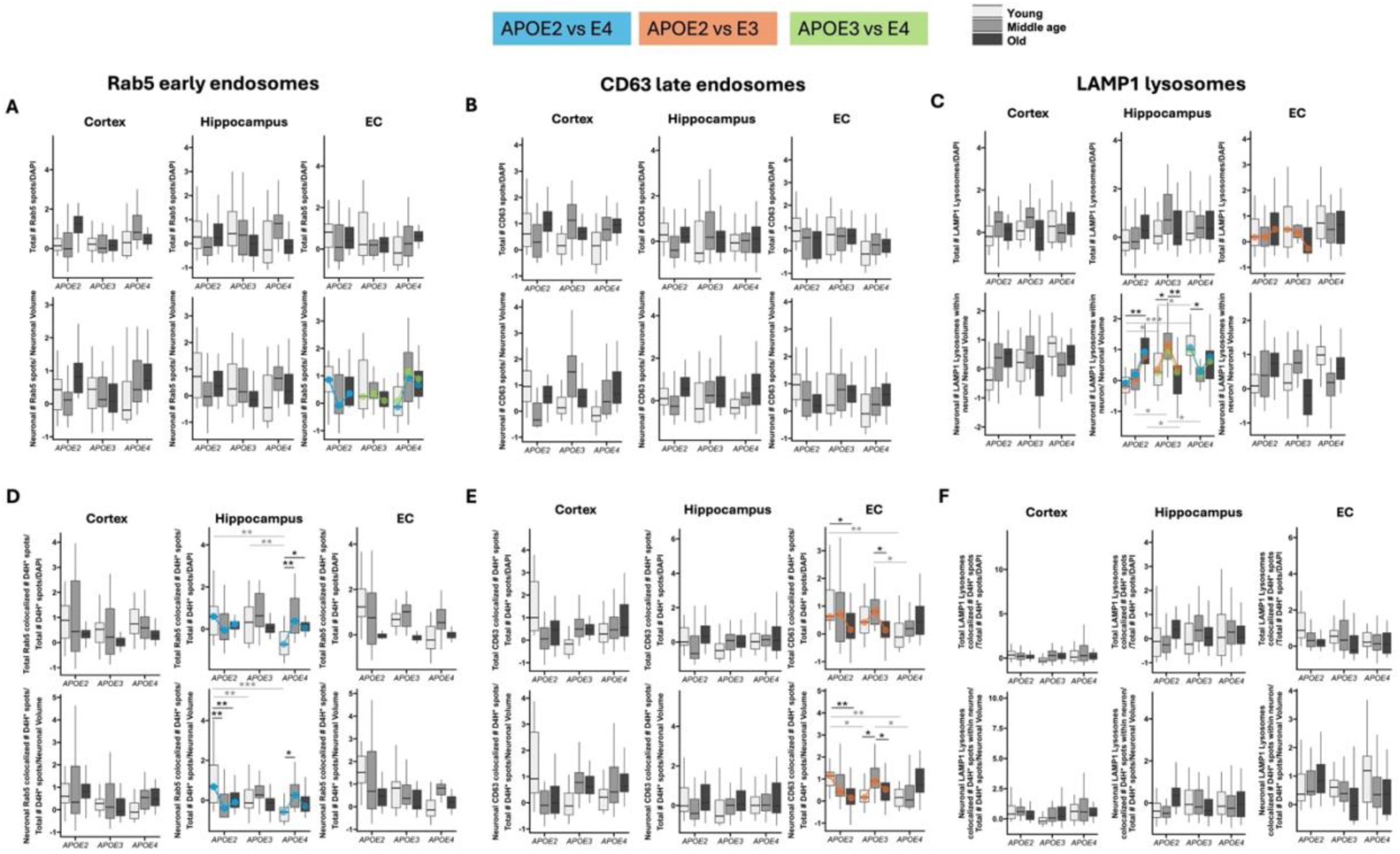
The interaction effect of age and *APOE* genotype on early endosomes (EE), late endosomes (LE), lysosomes (LY), and cholesterol associated with EE’s, LE’s, and LY’s. **A**) Box plots representing total number of Rab5 spots, normalized to number of DAPI nuclei. *Lower row*: box plots representing neuronal number of Rab5 spots normalized to the total NeuN neuronal volume. The same metrics but different markers are depicted in plots **B)**, and **C)**. **B)** CD63 spots, and **C)** Lamp1 particles. **D)** Box plots representing total Rab5 colocalized D4H* spots, normalized to the total number of D4H* spots and subsequently normalized to the number of DAPI nuclei. *Lower row*: box plots representing neuronal number of Rab5 colocalized D4H* spots, to normalized to the total number of D4H* spots, and subsequently to the total NeuN neuronal volume. The same metrics but different markers are depicted in plots **D)**, and **F)**. **E)** CD63 spots, and **F)** Lamp1 particles. The data is grouped by age; young, middle age, old. *Statistics:* data from individual experiments were Z-scored before pooling. Linear mixed-effects model was fitted for each variable using *sex*, *age*, and *APOE* genotype as fixed effects, and the interaction effect between *APOE* genotype and *age*. Estimated marginal means were then computed for the *APOE* × *Sex* interaction. Significant *age* x *APOE2* vs *APOE4* interaction effect depicted in blue overlay curve, *age* x *APOE2* vs *APOE3* in orange overlay curve, *age* x *APOE3* vs *APOE4* in green overlay curve. Significance bars in grey show *APOE* genotype pairwise comparisons, and in black show *age* dependent comparisons, p<0.05: *, p<0.01:**, p<0.001:***. N = 5 animals per sex, age, *APOE* genotype.

**Table 2.**
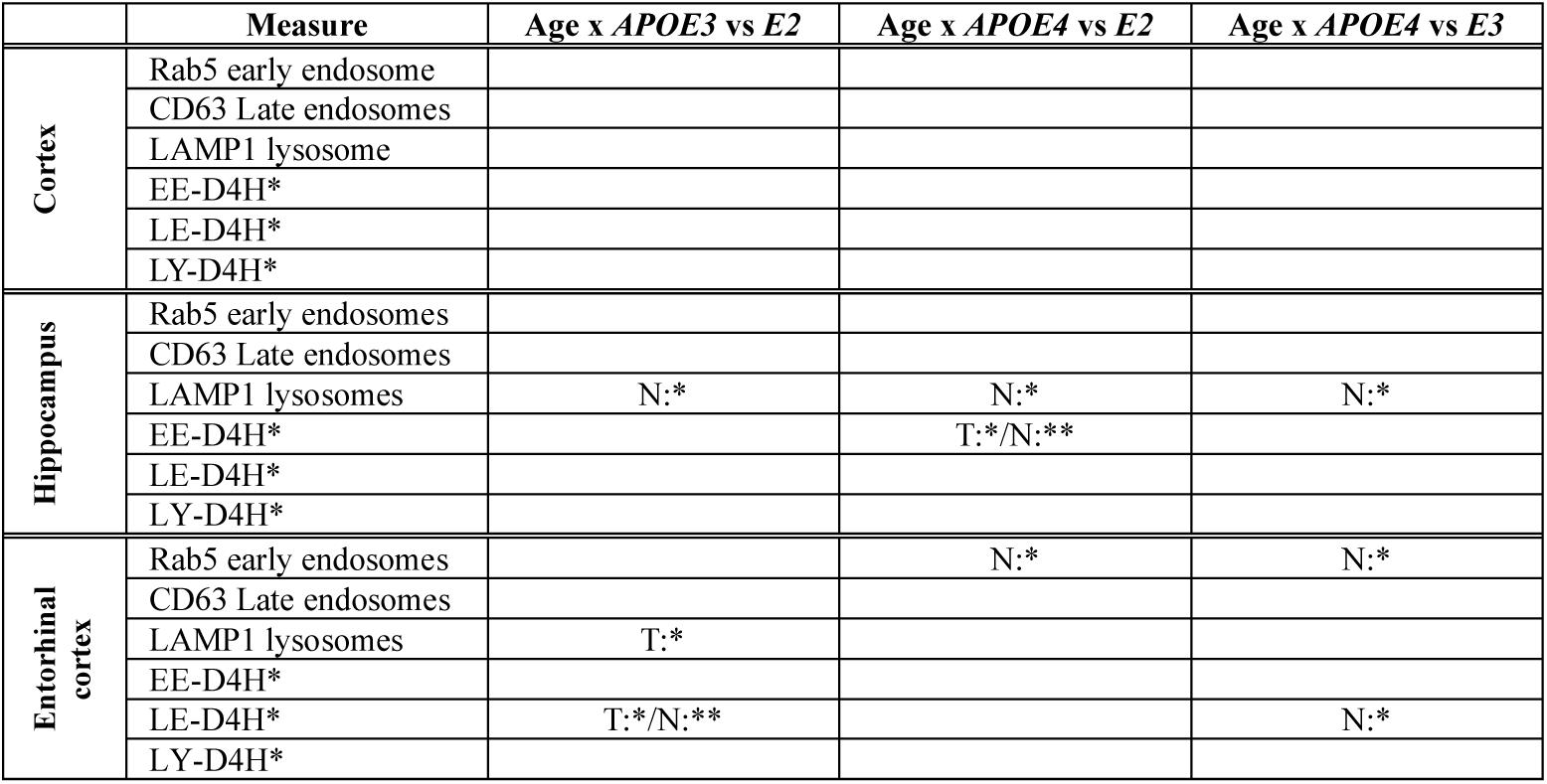
Interaction effects of *APOE* genotype x age on the endolysosomal system and endolysosomal cholesterol in the mouse brain.

The total and neuronal EE-Ch levels in the hippocampus showed a significant age effect dependent on the difference between *APOE2* and *APOE4* (**Fig. 3D**). This was primarily driven by an *APOE* dependent decrease at young age (*E2*>*E3*>*E4*). In total, EE-Ch levels showed a more prominent increase with age in *APOE4* (Y<M;Y<O), while neuronal levels showed a more prominent decrease with age in the *APOE2* genotype (Y>M;Y<O). These data indicate that EE-Ch levels are regulated by the interaction of *age x APOE*, exclusively in the hippocampus. Furthermore, both total and neuronal LE-Ch are significantly affected by the interaction effect of *age x APOE* in the EC (**Fig. 3E**). This is driven by the difference between *APOE2* and *APOE3* in LE-Ch levels with aging. The effect was slightly stronger in N-LE-Ch, driven by high levels at young age, declining with age in *APOE2* (Y>O), and a sharp peak at middle age in *APOE3* (Y<M>O). At the same time, at young age *APOE2>E3*; *E2>E4*, and at middle age *APOE3>E4*. Importantly, in the EC, cholesterol levels in the LE were affected by *age x APOE*, and in the hippocampus, EE-Ch was exclusively affected. There was no interaction effect on LY-Ch levels (**Fig. 3F**).

In the age effects plotted by *APOE* genotype, it was evident that there were prominent differences between genotypes in the young age group. To further investigate these *APOE* differences, the data from young mice only were fitted to a linear mixed-effects model, with *APOE* and sex as fixed effects. Pairwise contrasts were estimated between *APOE* genotypes, adjusted for sex (**Fig. 4**). In young animals, there was a striking pattern of EE, LE, and LY levels that were similar in each brain region (CTX, HIPP, and EC). In the CTX, there were no effects on total number of EE, or on total number of cholesterol colocalized with EE’s (**Fig. 4A, D**). However, there was a striking *APOE* allele dependent decrease in neuronal LE’s (*APOE2*>*E3*;*E2*>*E4*), while the opposite was observed for neuronal LY levels (*APOE2*<*E4*). At the same time, N-LE-Ch levels showed similar expression to N-LE’s, while N-LY-Ch levels did not show an *APOE* dependent change. Even though Rab5 Neuronal EE’s and N-EE-Ch were not significantly different, their levels showed a similar decreasing pattern with *APOE* genotype as N-LE and N-LE-Ch levels.

**Figure 4.**
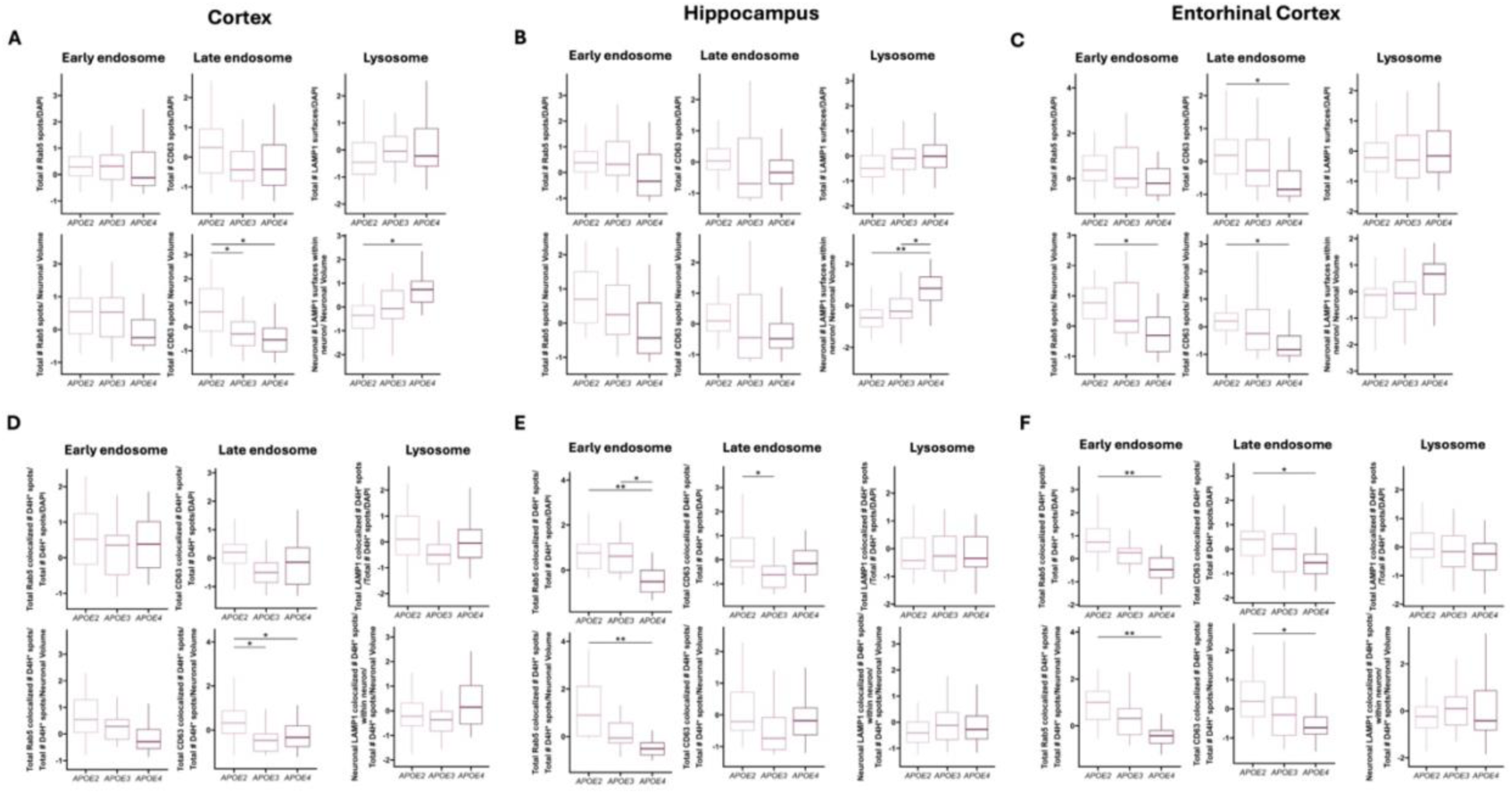
The effect of *APOE* genotype in young mice, in each brain region on early endosomes (EE), late endosomes (LE), lysosomes (LY), and cholesterol associated with EE’s, LE’s, and LY’s. **A**) Box plots representing total number of Rab5 spots (Early endosomes), normalized to number of DAPI nuclei, in the **cortex**. *Lower row*: box plots representing neuronal number of Rab5 spots normalized to the total NeuN neuronal volume. The same metrics but different markers are depicted in the middle plot; CD63 spots (Late endosomes), and far right plot; Lamp1 particles (Lysosomes). **B)** Hippocampus, **C)** Entorhinal cortex (EC), **D)** Box plots representing total Rab5 colocalized D4H* spots, normalized to the total number of D4H* spots and subsequently normalized to the number of DAPI nuclei, in the **cortex**. *Lower row*: box plots representing neuronal number of Rab5 colocalized D4H* spots, to normalized to the total number of D4H* spots, and subsequently to the total NeuN neuronal volume. The same metrics but different markers are depicted in the middle plot; CD63 spots (Late endosomes), and far right plot; Lamp1 particles (Lysosomes). **E)** Hippocampus, **F)** Entorhinal cortex (EC). The data is grouped by *APOE* genotype. *Statistics:* data from individual experiments were Z-scored. Linear mixed-effects model was fitted with *sex* and *APOE* genotype as fixed effects. Estimated marginal means were then computed for the *APOE*, and pairwise comparisons were measured between groups, p<0.05: *, p<0.01:**. N = 5 animals per sex, age, *APOE* genotype.

Altogether there was a clear neuron-specific effect in the young CTX, where the decrease in LE’s and LE-Ch in *APOE3* and *E4* may be overcompensated for by an increase in the number of LY’s in *APOE4* vs *APOE2*. However, the amount of cholesterol colocalized with LY’s was not increased in *APOE4* vs *APOE2*, indicating that there may be a failure of LY’s to process cholesterol in *APOE4* neurons.

In the HIPP, the total levels of EE’s, LE’s, and LY’s were not affected by *APOE* genotype (**Fig. 4B**). Furthermore, while both neuronal EE’s and LE’s showed an increasing pattern with *APOE* genotype, only N-LY’s showed a significant upregulation in *APOE4* (*APOE2*<*E4*;*E3*<*E4*). Both total and neuronal EE-Ch was reduced with *APOE4* (Total: *APOE2*>*E4*;*E3*>*E4*, neuronal: *APOE2*>*E4*), which shows that EE-Ch levels are sensitive to *APOE* genotype, but EE levels itself are not (**Fig. 4E**). There was also a decrease in T-LE-Ch in *APOE3*<*E2*, with no change in *APOE4*, and N-LE-Ch levels were not affected. Similar to the cortex, there was no effect of *APOE* genotype on total or neuronal LY-Ch, even though neuronal LY levels were increased, indicating that this upregulation is not linked to the LY capacity to process cholesterol.

In the EC, total EE levels were not affected, but there was a neuron-specific *APOE* effect on EE’s (*APOE2*>*E4*) (**Fig. 4C**). Both total and neuronal LE’s were decreased in *APOE4*<*E2*, while lysosomes were not affected by *APOE* genotype. There was a striking decrease in both total and neuronal EE-Ch and LE-Ch (*APOE4*<*E2*), while there was a complete absence of *APOE* effect on LY-Ch levels (**Fig. 4F**).

Altogether, there was a clear involvement of reduced EE-Ch levels in both the HIPP and EC of the young mice the *APOE4* genotype. Further, a reduction in total and neuronal LE’s and LE-Ch was prominent in the EC, while only neuron-specific LE and LE-Ch were decreased in the cortex, with this effect absent in the HIPP. Lastly, only neuron-specific LY levels were increased with *APOE* genotype in the CTX, but more significantly in the HIPP. There was no regulation of total or neuronal LY-Ch levels in any of the brain regions. This indicates that with reduced EE and LE levels in E4<E3<E2 of the young mice, there may be a reduced capacity of EE’s in HIPP and EC, as well as LE’s in the CTX and EC to process cholesterol. While there was a neuronal LY increase in HIPP and CTX, this was not paired with an increase in cholesterol colocalization, indicating that neuronal *APOE4* lysosomes in young brains may have reduced cholesterol processing capacity.

*Interestingly, these striking APOE genotype effects were not observed at middle age or old age* (**Fig. S3** and **Fig. S4**). However, only in the EC of old animals, there was a small decrease of neuronal LY’s in *APOE3* and an increase in *APOE4* (*APOE2*>*E3*<E4) (**Fig. S4C, F**). N-LY-Ch levels in the HIPP and EC were also decreased in *APOE3<APOE2*; and a decrease in *APOE4*<*APOE2* (EC only). This effect could indicate that in old age, in neurons in the EC, *APOE4* lysosomes increase (*E3*<*E4*), but the LY capacity to process cholesterol does not increase, showing that this effect observed in young animals, is also present at old age, specifically in the EC.

### Effect of sex and *APOE* genotype and aging on the endolysosomal system and endolysosomal cholesterol

Finally, we also performed sex-specific analyses on all of our data, given that *APOE4* penetrance is known to be strongly affected by sex, with female *APOE4* carriers being at an increased risk of AD compared to male *APOE4* carriers^3^. There was only one significant main effect of sex, and this was on T-LY’s in the EC (**Table 1**). There was not a pairwise significance comparing male and female mice (adjusted for *APOE* and *age*) on T-LY levels, but T-LY-Ch and N-LY-Ch in the EC were significantly increased in females, indicating that lysosomes in the EC in female mice contain more cholesterol (**Fig. 5**).

**Figure 5.**
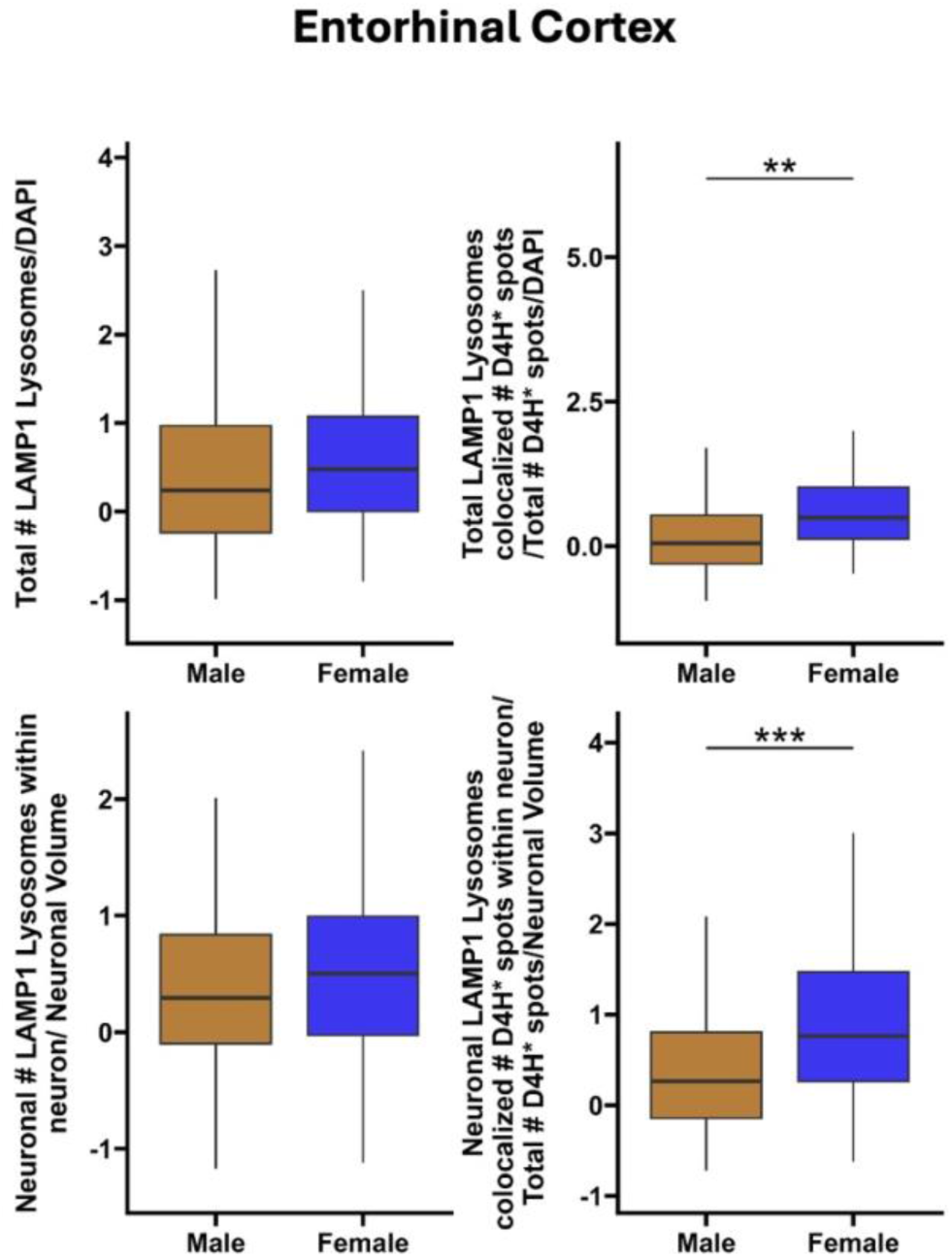
The main effect of sex on lysosomes (LY) and LY-cholesterol in the EC. Box plots representing total number of Lamp1 particles (lysosomes), normalized to number of DAPI nuclei, in the EC. *Lower row*: box plots representing neuronal number of Rab5 spots normalized to the total NeuN neuronal volume. Data grouped by sex: male (ochre fill) and female (blue fill). *Statistics:* data from individual experiments were Z-scored. Linear mixed-effects model was fitted with *sex*, *APOE* genotype, and *age* as fixed effects. Estimated marginal means were then computed for the *sex*, and pairwise comparisons were measured between groups, p<0.05: *, p<0.01:**, p<0.001:***. N = 5 animals per sex, age, *APOE* genotype.

We also observed *sex* x *age* interaction effects and *sex* x *APOE* genotype interaction effects (**Table 3**). The majority of *sex* x *age* interactions were observed in the EC (**Fig. 6**), with one exception - N-LY’s in the HIPP showed a significant *sex* x *age* interaction effect (**Fig. S5**). This effect in the HIPP was mainly driven by the increase in N-LY’s in females vs males at old age. There were no significant interaction effects on EE or LE particles in the EC. However, there was a highly significant effect on T-LY’s in the EC, driven by an age-related increase in T-LY’s in females, and an age-related decrease in males (**Fig. 6A**). This effect also showed a pairwise difference in the young group, where males had higher T-LY levels than females, in contrast to old age, where females had increased T-LY levels compared to males. Furthermore T-LY levels were significantly lower in old males versus young males. No further effects were observed in the HIPP and CTX (**Fig. S5A, C**).

**Figure 6.**
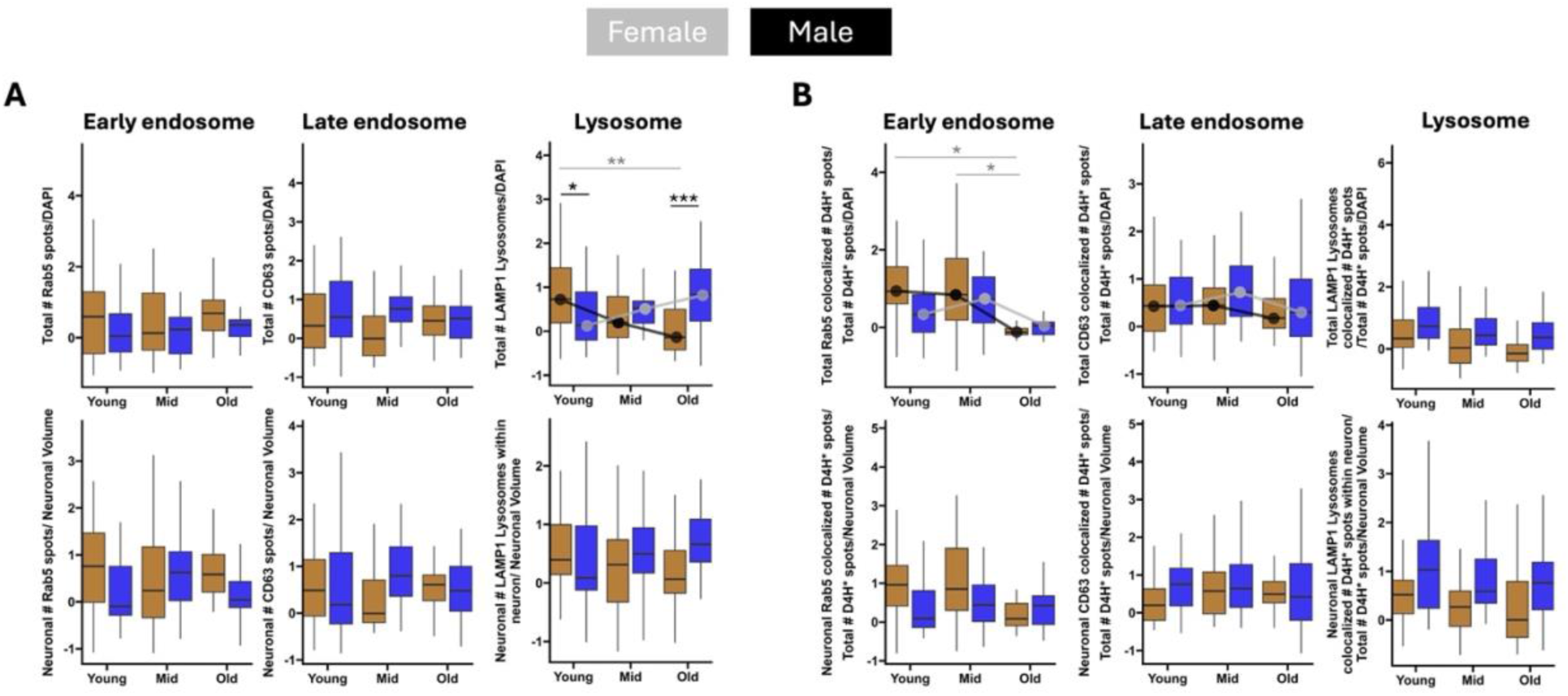
The interaction effect of sex x age in the EC on early endosomes (EE), late endosomes (LE), lysosomes (LY), and cholesterol associated with EE’s, LE’s, and LY’s. **A)** Box plots representing total number of Rab5 spots, CD63 spots, and LAMP1 particles normalized to number of DAPI nuclei. *Lower row*: box plots representing neuronal number of Rab5 spots, CD63 spots, and LAMP1 particles normalized to the total NeuN neuronal volume. The same metrics but different markers are depicted in plots **B)** Box plots representing total Rab5, CD63 and LAMP1 colocalized D4H* spots, normalized to the total number of D4H* spots and subsequently normalized to the number of DAPI nuclei. *Lower row*: box plots representing neuronal number of Rab5, CD63, and LAMP1 colocalized D4H* spots, to normalized to the total number of D4H* spots, and subsequently to the total NeuN neuronal volume. Data was grouped by age; young, middle age, old, and sex; male (ochre fill) and female (blue fill). *Statistics:* data from individual experiments were Z-scored. Linear mixed-effects model was fitted for each variable using *sex*, *age*, and *APOE* genotype as fixed effects, and the interaction effect between *sex* and *age*. Estimated marginal means were then computed for the *age* × *Sex* interaction. Significant *age* x sex interaction effect depicted in grey overlay curve (female), and black overlay curve (male). Significance bars in grey show *age* pairwise comparisons, and in black show *sex* dependent comparisons, p<0.05: *, p<0.01:**, p<0.001:***. N = 5 animals per sex, age, *APOE* genotype.

**Table 3.** Interaction effects of sex x *APOE* genotype x age on the endolysosomal system and cholesterol levels in the brain.

|  | Measure | Age x Sex interaction effect | *APOE* x Sex effect | Sex x *APOE* x Age<br>(*APOE3*) | Sex x *APOE* x Age<br>(*APOE4*) |
| --- | --- | --- | --- | --- | --- |
| Cortex | Rab5 early endosome |  |  |  |  |
|  | CD63 Late endosomes |  |  |  | T:* |
|  | LAMP1 lysosome |  |  |  |  |
|  | EE-D4H* |  |  |  |  |
|  | LE-D4H* |  |  |  |  |
|  | LY-D4H* |  |  |  |  |
| Hippocampus | Rab5 early endosome |  |  |  |  |
|  | CD63 Late endosomes |  |  |  |  |
|  | LAMP1 lysosome | N:* |  |  |  |
|  | EE-D4H* |  | N:* |  |  |
|  | LE-D4H* |  |  |  |  |
|  | LY-D4H* |  |  |  |  |
| Entorhinal cortex | Rab5 early endosome |  |  |  |  |
|  | CD63 Late endosomes |  |  |  |  |
|  | LAMP1 lysosome | T:*** | T:* | T:** |  |
|  | EE-D4H* | T:* |  |  |  |
|  | LE-D4H* | T:* |  | T:* | T/N:* |
|  | LY-D4H* |  |  |  |  |

Cholesterol associated with EE’s and LE’s were affected by the sex x age interaction in the EC, while this effect was absent for LY-Ch (**Fig. 6B**). For T-EE-Ch, the effect was driven by a decrease in old males (Y>M;M>O), while EE-Ch levels were also reduced in old females, though this was not significant. The T-EE-Ch levels in males fit a steady downward slope with age, while in females, T-EE-Ch levels only dropped off in old age. Similar slopes were observed in T-LE-Ch levels, but there were no pairwise differences. No further effects were detected in the HIPP and CTX (**Fig. S5B, D**).

These data indicate that the EC is particularly sensitive to interaction effects of *sex* x *age*. Both total and neuronal LY levels were increased in old females compared to males, but sex x age did not affect lysosomal cholesterol. This is contrary to EE and LE particles in the EC, which were not affected, while total EE-Ch and LE-Ch levels followed a different trajectory with age in females or males.

The second interaction term that was tested was *sex x APOE* (**Fig. 7**). Levels of LE’s or LE-Ch were not affected. There were only two significant effects, one of which was a significant effect on neuronal EE-Ch in the HIPP (**Fig. 7B**). EE particles were not affected (**Fig. 7B**), nor were the EE-Ch levels affected in any other brain region. N-EE-Ch levels in HIPP were the same in all *APOE* genotypes in females, while in males there was a slight increase in *APOE3*, and a decrease in *APOE4*. Furthermore, while there was no interaction effect for N-EE-Ch levels, pairwise contrasts did show a significant decrease between females and males carrying the *APOE3* genotype, while there was no contrast between sexes for *APOE2*, nor *APOE4* (**Fig. 7B**). Male and female *APOE2* mice possessed increased N-EE-Ch compared to their *APOE4* and *APOE3* counterparts, respectively. This indicates that neurons in the EC of female brains carrying *APOE3*, vs males, have lower levels of cholesterol in EE’s, implying lower levels of cholesterol uptake. The levels of N-EE-Ch in *APOE3* females was similar to those observed in *APOE4* males and females.

**Figure 7.**
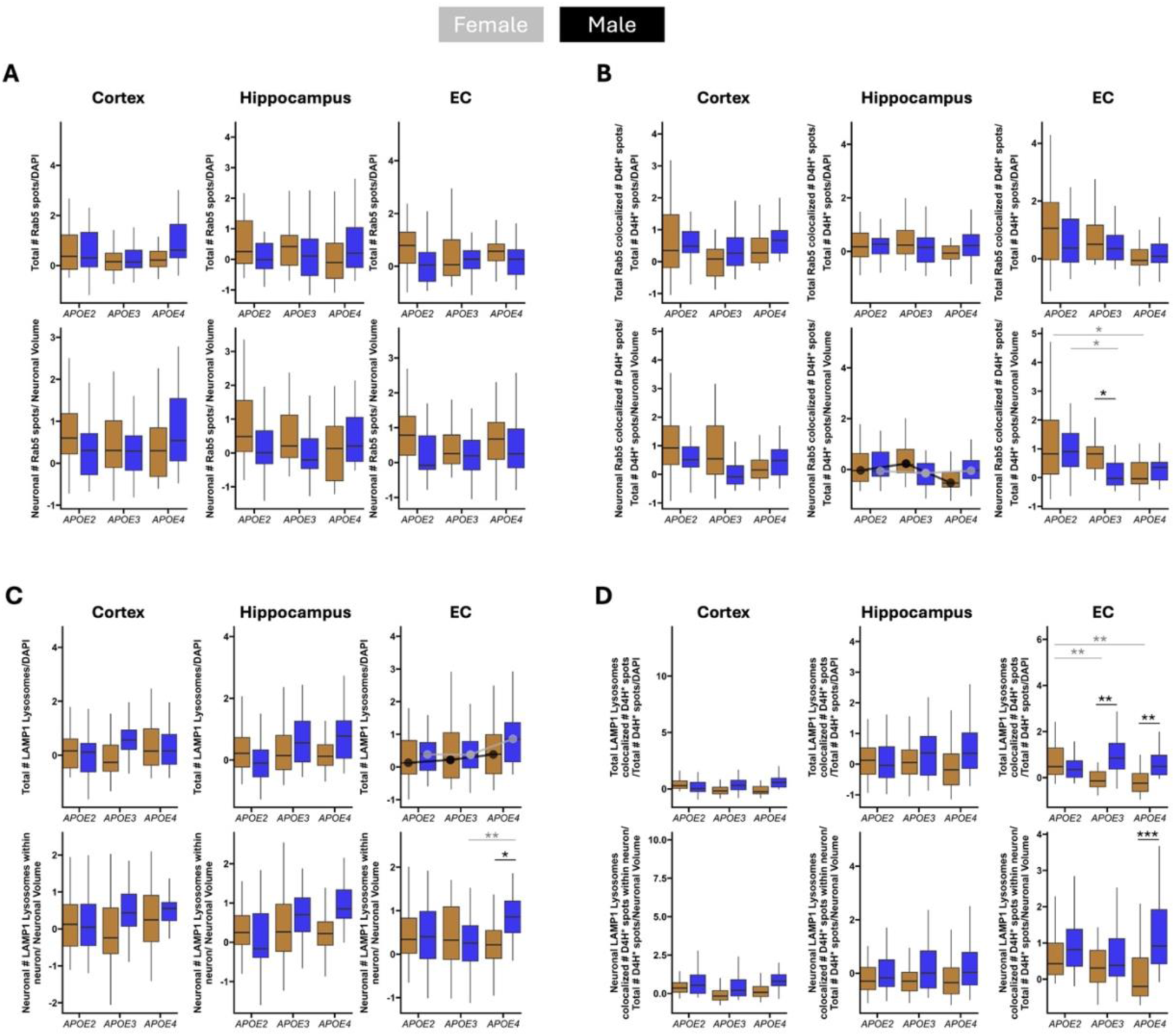
The interaction effect of sex x *APOE* genotype early endosomes (EE), EE-cholesterol, lysosomes (LY) and LY-cholesterol. **A)** Box plots representing total number of Rab5 spots and **C)** Lamp1 lysosomal particles normalized to number of DAPI nuclei in the cortex, hippocampus, and entorhinal cortex (EC). *Lower row*: box plots representing neuronal number of Rab5 spots and Lamp1 lysosomal particles normalized to the total NeuN neuronal volume in the cortex, hippocampus, and entorhinal cortex (EC). **B)** Box plots representing total Rab5 and **D)** Lamp1 lysosomal particles colocalized D4H* spots, normalized to the total number of D4H* spots and subsequently normalized to the number of DAPI nuclei in the cortex, hippocampus, and entorhinal cortex (EC). *Lower row*: box plots representing neuronal number of Rab5 and Lamp1 lysosomal particles colocalized D4H* spots, to normalized to the total number of D4H* spots, and subsequently to the total NeuN neuronal volume in the cortex, hippocampus, and entorhinal cortex (EC). Data was grouped by *APOE* genotype; *APOE2*, *APOE3*, *APOE4*, and sex; male (ochre fill) and female (blue fill). *Statistics:* data from individual experiments were Z-scored. Linear mixed-effects model was fitted for each variable using *sex*, *age*, and *APOE* genotype as fixed effects, and the interaction effect between *sex* and *APOE*. Estimated marginal means were then computed for the *APOE* × *Sex* interaction. Significant *APOE* x sex interaction effect depicted in grey overlay curve (female), and black overlay curve (male). Significance bars in grey show *APOE* genotype pairwise comparisons, and in black show *sex* dependent comparisons, p<0.05: *, p<0.01:**, p<0.001:***. N = 5 animals per sex, age, *APOE* genotype.

Total levels of LAMP1 LY particles were also affected by sex, dependent on *APOE* genotype in the EC region (**Fig. 7C**). This was driven by a sharper incline in LY’s in females comparing *APOE4* to *APOE3*, versus males. There were no further *sex* x *APOE* main factor interaction effects in other brain regions. However, paired contrasts showed that N-LY’s in the EC were increased in *APOE4* female compared to male mice, and compared to *APOE3* females. These results indicate that, specifically *APOE4 females are vulnerable to increased levels of total and N-LY’s in the EC*. The N-LY levels in the HIPP was reminiscent of this effect, but not significant. Furthermore, T-LY-Ch was decreased in males by genotype (*APOE3*<*E2*;*E4*<*E2*) in the EC, and females had increased levels of T-LY-Ch in the *APOE3* or *APOE4* genotype compared to their male counterparts. Finally, specifically N-LY-Ch levels were only increased in females vs males with *APOE4* genotype. These results indicate that there are striking differences in total and neuronal LY’s, and total and neuronal LY-Ch, in males and females across *APOE* genotypes. Thus, females carrying the *APOE4* genotype were more susceptible than males to an increase in N-LY-Ch in the EC, suggesting cholesterol accumulation.

Neither CD63 late endosomes, nor LE-Ch particles were affected by a *sex x APOE* interaction effect (**Fig. S3**).

Next, a three-way interaction effect of *sex x Age x APOE* was tested by fitting a linear mixed-effects model using *sex*, *APOE* genotype, and *age* as fixed effects, with a random intercept for image ID to account for repeated measures (**Table 3**). This was done for the cortex, hippocampus, and EC.

There were no significant three-way interactions for Rab5 EE’s, or EE-Ch (**Fig. 8**). Even though there were no interactions of the main variables, there were significant pairwise differences between *APOE* genotypes or *sex* per age group. In the cortex, total levels of EE’s were upregulated in *APOE4* males versus *APOE2* (**Fig. 8A**). While at old age, there was a decrease in T-EE’s in *APOE3* and *APOE4* females compared to *APOE2*. Interestingly, N-EE levels at old age were upregulated in *APOE4* females compared to *APOE3*, pointing to a neuron-and female-specific upregulation in *APOE4*. In total EE-Ch levels, there was a striking decrease in *APOE3* males versus females, while at old age there was no sex-dependent difference, but *APOE3* females had decrease EE-Ch levels compared to both *APOE2* and *APOE4* females (**Fig. 8B**). In neuronal EE-Ch, a similar *APOE3* female decrease was observed, which in this case was also decreased compared to male *APOE3*.

**Figure 8.**
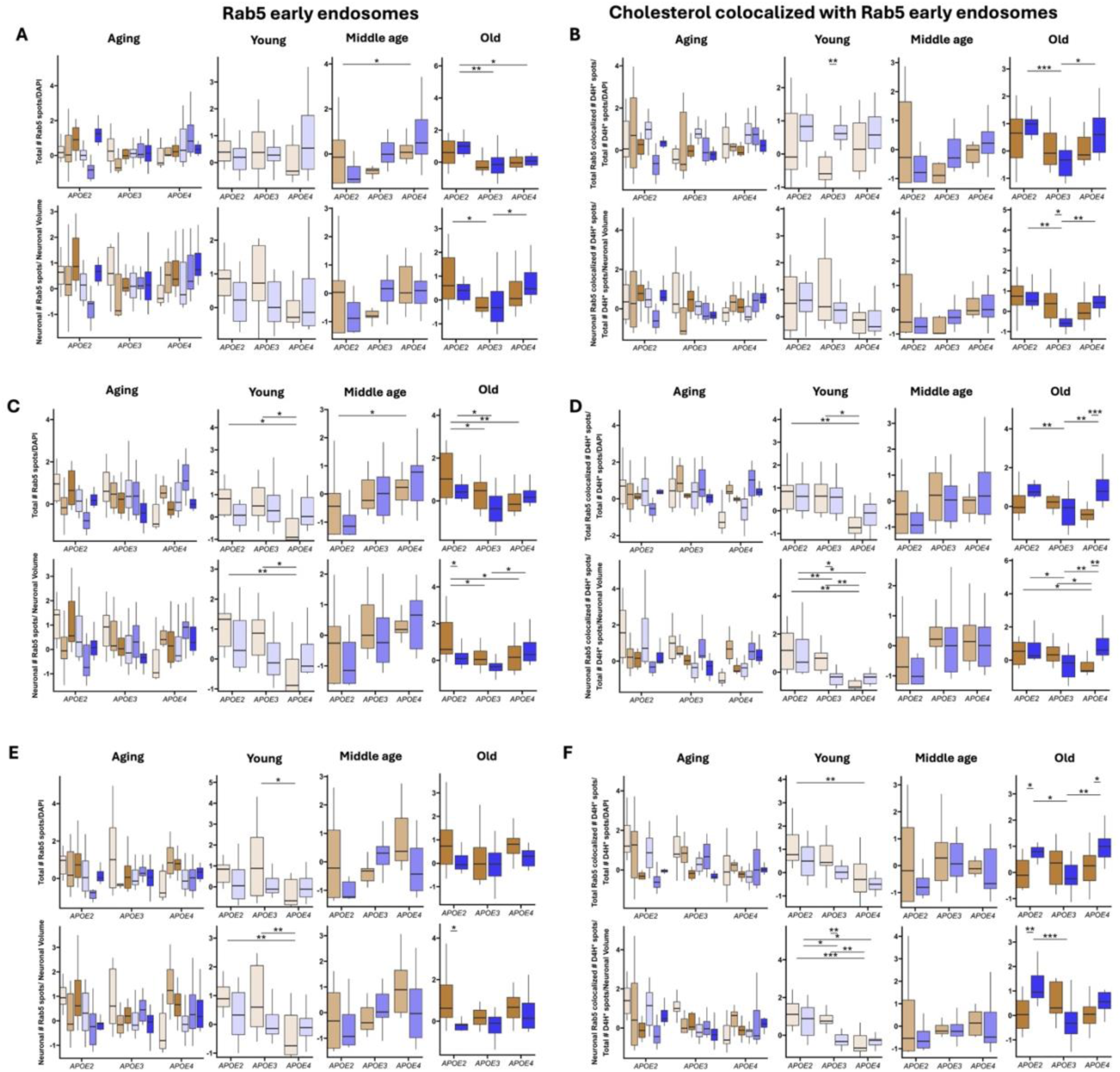
The three-way interaction effect of sex x *APOE* genotype x age on early endosomes (EE), EE-cholesterol. **A)** Box plots representing total number of Rab5 spots normalized to number of DAPI nuclei throughout aging, at young, middle, and old age. *Lower row*: box plots representing neuronal number of Rab5 spots and normalized to the total NeuN neuronal volume throughout aging, at young, middle, and old age. **B)** Box plots representing total Rab5 and colocalized D4H* spots, normalized to the total number of D4H* spots and subsequently normalized to the number of DAPI nuclei in the cortex, hippocampus, and entorhinal cortex (EC). *Lower row*: box plots representing neuronal number of Rab5 colocalized D4H* spots, to normalized to the total number of D4H* spots, and subsequently to the total NeuN neuronal volume throughout aging, at young, middle, and old age. **A)** and **B)** cortex, **C)** and **D)** hippocampus, **E)** and **F)** entorhinal cortex (EC) Data was grouped by *APOE* genotype; *APOE2*, *APOE3*, *APOE4*, sex; male (ochre fill) and female (blue fill), and age; young (alpha 0.2), middle age (alpha 0.6), old (alpha 1). *Statistics:* data from individual experiments were Z-scored. For the “Aging” plot, linear mixed-effects model was fitted for each variable using *sex*, *age*, and *APOE* genotype as fixed effect. Wald χ² tests (HC3-robust) evaluated three-way *sex*×*APOE*×*age* interaction effects. For the individual plots within age group young, middle age and old, linear mixed-effects model was fitted with *sex* and *APOE* genotype as fixed effects. Wald χ² tests (HC3-robust) evaluated the interaction effect of *sex*×*APOE*, male within *APOE* comparisons, female within *APOE* comparisons, *sex* differences within each *APOE* genotype. P-values were FDR-corrected across contrasts, p<0.05: *, p<0.01:**, p<0.001:***.. N = 5 animals per sex, age, *APOE* genotype.

In the hippocampus there were numerous pairwise interactions. At young age, there was a male-specific decrease of *APOE4* T-EE levels compared to *APOE2* and *APOE3* (**Fig. 8C**). Whereas, there was a significant increase in *APOE4* males versus *APOE2* at middle age, which was followed again by a decrease in old age. Here, *APOE2* males had the highest T-EE levels compared to *APOE3* and *APOE4*, while female *APOE2* T-EE levels were increased only compared to *APOE3*, and no further difference was observed with *APOE4*. Neuron-specific EE levels in old hippocampus were the same as T-EE levels, with a decrease in *APOE3* and *APOE4* males compared to *APOE2* males. No significant differences at middle age were observed, and N-EE levels showed an *APOE2* male-specific increase compared to male *APOE3*, male *APOE4*, and female *APOE3* in old age. Interestingly, N-EE levels were increased in *APOE4* females compared to *APOE3* at old age, which was not observed for T-EE-Ch levels.

Next, total EE-Ch levels were specifically decreased in *APOE4* males compared to *APOE2* and *APOE3* males at young age (**Fig. 8D**). No differences were observed at middle age, and a decrease in *APOE3* versus *APOE2* females was observed, followed by an increase in *APOE4* females, compared to *APOE3* females, and *APOE4* males, at old age. This resulted in an *APOE4* female-specific increase in T-EE-Ch at old age in the hippocampus.

Lastly, neuron-specific EE-Ch showed a male *APOE4*-specific decrease compared to *APOE2* and *APOE3*. On the other hand, there was a female-specific upregulation observed in *APOE2* females compared to *APOE3* and *APOE4* females. Consequently, *APOE3* females had lower N-EE-Ch levels than *APOE3* males. This demonstrates that at young age, *APOE3* males have N-EE-Ch levels similar to *APOE2* males, and *APOE3* females have levels similar to *APOE4* females in the hippocampus. At old age, the male-specific decrease in *APOE4* (vs. *APOE3* and *APOE2*) was still observed. On the other hand, while *APOE3* females still had lower N-EE-Ch levels than *APOE2*, *APOE4* females had increased levels compared both to *APOE3* female and *APOE4* male. These results demonstrate that there is a female-specific increase in total and neuronal cholesterol in the EE in old age, indicating more early endosomal cholesterol uptake, which may be functionally linked to the decreased EE-Ch levels in *APOE4* females, but only in neurons.

There were fewer significant pairwise comparisons of EE and EE-Ch levels in the EC at old age, but changes at young age in the EC were largely equivalent to the hippocampus (**Fig. 8**). At young age, there was a significant decrease in *APOE4* males compared to *APOE3*, and no further significant changes in middle and old age (**Fig. 8E**). The same decrease can was observed in N-EE levels at young age, with the added significant decrease in *APOE4* males and *APOE2*. Furthermore, T-EE-Ch levels were significantly decrease in *APOE4* males versus *APOE2* (**Fig. 8F**). No further differences were observed in middle-aged mice, and in the old mice, there was a female *APOE2*-specific increase in N-EE-Ch compared to *APOE3*, and male *APOE2*. On the other hand, *APOE4* females had increased levels compared to *APOE3*, and male *APOE4*. Lastly, neuron-specific EE-Ch levels at young age had the same levels as in the EC, *APOE2* male and female, and *APOE3* male were increased compared to *APOE3* female, and *APOE4* male and female. At old age, there was an increase in N-EE-Ch in *APOE2* females compared to *APOE3* and *APOE2* males.

Taken together there is effectively a consistent upregulation of total and neuronal EE-Ch in *APOE4* females at old age, in all three brain regions, indicating that this effect is not specific to the medial temporal lobe. Furthermore, the hippocampus and EC showed very similar levels of EE’s and EE-Ch, especially for N-EE-Ch levels in young brains, indicating that EE functions and cholesterol uptake are functionally linked between these regions.

In contrast, CD63 total LE levels in the cortex were significantly different between *APOE4* and *APOE2*, dependent on *sex* and *age* (**Fig. 9**). The three-way interaction effect of *age* x *sex* x *APOE* was significant, and the interaction curves showed a sharp V (in LE levels) in male *APOE2* carriers with age, while female *APOE2* carriers showed steady levels of LE’s with aging (**Fig. 9A**). On the other hand, LE levels in *APOE4* males showed relatively steady levels with aging, whereas the females showed a sharp increase from young to middle age, and steady levels to old age. There were no further significant effects dependent on *APOE3*.

**Figure 9.**
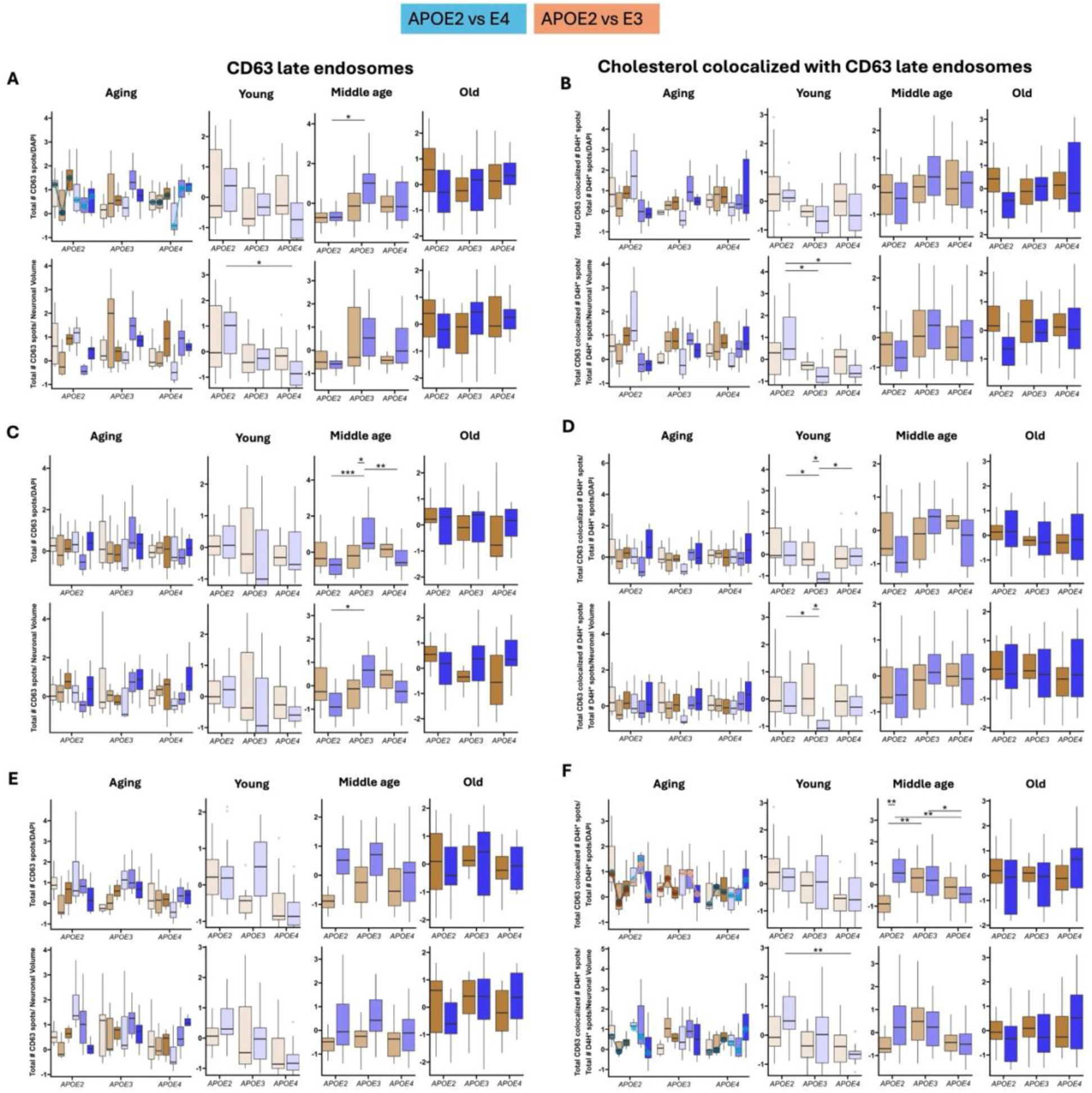
The three-way interaction effect of sex x *APOE* genotype x age on late endosomes (LE), LE-cholesterol. **A)** Box plots representing total number of CD63 spots normalized to number of DAPI nuclei throughout aging, at young, middle, and old age. *Lower row*: box plots representing neuronal number of CD63 spots normalized to the total NeuN neuronal volume throughout aging, at young, middle, and old age. **B)** Box plots representing total CD63 and colocalized D4H* spots, normalized to the total number of D4H* spots and subsequently normalized to the number of DAPI nuclei in the cortex, hippocampus, and entorhinal cortex (EC). *Lower row*: box plots representing neuronal number of CD63 colocalized D4H* spots, to normalized to the total number of D4H* spots, and subsequently to the total NeuN neuronal volume throughout aging, at young, middle, and old age. **A)** and **B)** cortex, **C)** and **D)** hippocampus, **E)** and **F)** entorhinal cortex (EC) Data was grouped by *APOE* genotype; *APOE2*, *APOE3*, *APOE4*, sex; male (ochre fill) and female (blue fill), and age; young (alpha 0.2), middle age (alpha 0.6), old (alpha 1). *Statistics:* data from individual experiments were Z-scored. For the “Aging” plot, linear mixed-effects model was fitted for each variable using *sex*, *age*, and *APOE* genotype as fixed effect. Wald χ² tests (HC3-robust) evaluated three-way *sex*×*APOE*×*age* interaction effects. For the individual plots within age group young, middle age and old, linear mixed-effects model was fitted with *sex* and *APOE* genotype as fixed effects. Wald χ² tests (HC3-robust) evaluated the interaction effect of *sex*×*APOE*, male within *APOE* comparisons, female within *APOE* comparisons, *sex* differences within each *APOE* genotype. P-values were FDR-corrected across contrasts, p<0.05: *, p<0.01:**, p<0.001:***.. N = 5 animals per sex, age, *APOE* genotype.

Zooming in on the effect of *sex x APOE* by *age* (Young, Middle age, Old), pairwise contrasts did not show any significant effect at young age, even though there was a decreasing trend in LE levels in females (*E2*<*E3*<*E4*), and at middle age a significant increase in *APOE3* vs *APOE2* was observed in females. Pairwise tests did not show any effects in the males, and no further differences in old age. While there was no interaction effect observed in neuron-specific LE levels (**Fig. 9A** bottom row), pairwise contrasts between sexes and *APOE* genotype per age group showed a decrease in N-LE levels in young *APOE4* females versus *APOE2*. There were no further pairwise differences at middle and old age. Total and neuronal LE-Ch levels were not affected by three-way interaction effects (**Fig. 9B**). There were no significant pairwise contrasts in T-LE-Ch levels, while N-LE-Ch levels were decreased in *APOE3* and *APOE4* females compared to *APOE2* at young age in the cortex. Altogether, the *sex* dependent effect on *age* that was different between *APOE2* and *APOE4* on total LE levels in the cortex demonstrates that there is an *APOE4* female-specific incline with aging. This pattern can be observed as a trend in the other metrics of LE levels in the cortex. Therefore, while neuronal LE’s and N-LE-Ch were decreased in *APOE4* females in young age, these levels in the cortex were affected throughout life.

In the hippocampus, no three-way interaction effects were detected. However, some pairwise differences were observed at young and middle age (**Fig. 9C-D**). Total LE levels were specifically increased at middle age in *APOE3* females, compared to both *APOE2* and *APOE4* females, and *APOE3* males (**Fig. 9C**). While no further comparisons were observed in old age. In N-LE levels specifically, there was also an increase in *APOE3* females detected, but this was only significant compared to *APOE2* females. Interestingly, LE-cholesterol levels only showed differences at young age in the hippocampus (**Fig. 9D**). Total LE-Ch was decreased in *APOE3* females compared to *APOE2* and *APOE4*, as well as *APOE3* males. Meanwhile, a similar effect, though less pronounced, was observed for N-LE-Ch levels; a decrease in *APOE3* females compared to *APOE2*, and *APOE3* males. These results indicate that there may be a functional relation of opposing regulation between LE-Ch in young females and LE particles in middle-aged females, where the effect on total levels was more pronounced.

In the EC, there were no significant three-way interactions observed affecting CD63 LE levels, and no pairwise differences were detected (**Fig. 9E**). Notably, total LE-cholesterol levels were significantly affected by the interaction of *sex* and *age* dependent on the difference between *APOE2* vs *APOE3*, and *APOE2* vs *APOE4* (**Fig. 9F**). In male *APOE2* carriers, T-LE-Ch levels follow a V pattern with aging, while in *APOE3* males, these followed a shallow peak profile. In *APOE2* females, T-LE-Ch levels rose sharply from young to middle age and showed a sharp drop at old age. The *APOE3* females showed steady levels from young to middle age, with a sharp drop in levels at old age. *APOE4* males showed similar T-LE-Ch levels with aging compared to *APOE3*, albeit with a sharper increase from young to middle age initially. Interestingly, the T-LE-Ch levels in *APOE4* females followed an inverse profile relative to the *APOE3* females: steady levels between young ang middle age, followed by a sharp increase at old age. Pairwise difference at young age showed a declining trend of T-LE-Ch with *APOE* genotype (*E2*>*E3*>*E4*) in the EC, which was significant in neuron-specific LE-Ch levels in females. At middle age, T-LE-Ch levels were significantly increased in females vs. males of the *APOE2* mice, and *APOE3* males were also increased, respectively. Furthermore, there was a decrease in *APOE4* females compared to *APOE2* and *APOE3*, but there were no further differences in T-LE-Ch levels in *APOE4* males. This suggests a female *APOE2*-specific increase and a female *APOE4*-specific decrease in T-LE-Ch at middle age in the EC. There were no further pairwise contrasts in the old age group.

The neuron-specific LE-Ch levels were only affected by the interaction of *sex* and *age* dependent on *APOE2* vs *APOE4* (**Fig. 9F** bottom row). This was marked in *APOE2* males by the decline of N-LE-Ch levels at mid age, and a slight increase at old age, while *APOE4* males showed a steady increase with aging. *APOE2* females showed a sharp decline of N-LE-Ch levels with aging, whereas *APOE4* females showed a decrease at middle age, and a sharp increase at old age. These results demonstrate that N-LE-Ch levels do not follow a simple decline or incline over age, but show distinct patterns dependent on *APOE* genotype and sex, during aging in the EC. In the young age group, N-LE-Ch levels were significantly higher in *APOE2* females compared to *APOE4*, in pairwise comparisons. There were no further differences at middle and old age. These results demonstrate that changes at young age are neuron-specific, while middle age changes in LE-Ch can only be observed in total levels, implying that cholesterol trafficking is regulated based on age and cell-type.

There were no three-way interactions on LAMP1 LY levels in the cortex (**Fig. 10**). Total-LY levels at old age in *APOE3* males were significantly decreased compared to *APOE4* and female *APOE3*, as shown with pairwise comparison testing (**Fig. 10A**). Neuronal-LY levels were significantly increased in young *APOE4* females, compared to *APOE2*, and N-LY levels at old age showed similar regulation to T-LY levels, with the addition of a significant difference between *APOE2* and *APOE3* in males. The only significant difference measured on N-LY-Ch levels was in the *APOE4* females, which showed a significant increase compared to *APOE2* and *APOE3* females, and *APOE4* males in the cortex (**Fig. 10B**).

**Figure 10.**
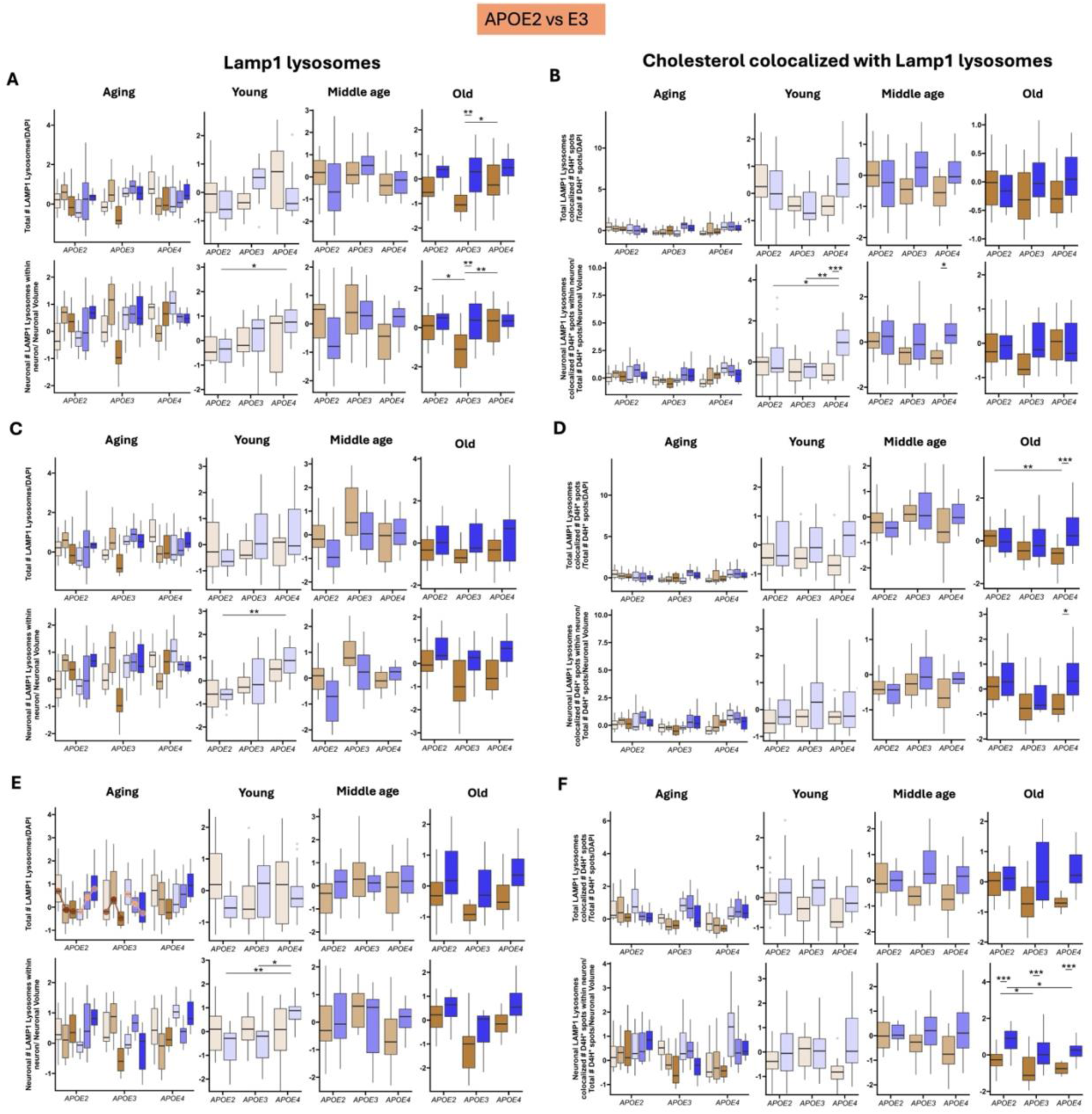
The three-way interaction effect of sex x *APOE* genotype x age on lysosomes (LY), LY-cholesterol. **A)** Box plots representing total number of Lamp1 particles normalized to number of DAPI nuclei throughout aging, at young, middle, and old age. *Lower row*: box plots representing neuronal number of Lamp1 particles normalized to the total NeuN neuronal volume throughout aging, at young, middle, and old age. **B)** Box plots representing total Lamp1 and colocalized D4H* particles, normalized to the total number of D4H* spots and subsequently normalized to the number of DAPI nuclei in the cortex, hippocampus, and entorhinal cortex (EC). *Lower row*: box plots representing neuronal number of Lamp1 colocalized D4H* particles, to normalized to the total number of D4H* spots, and subsequently to the total NeuN neuronal volume throughout aging, at young, middle, and old age. **A)** and **B)** cortex, **C)** and **D)** hippocampus, **E)** and **F)** entorhinal cortex (EC) Data was grouped by *APOE* genotype; *APOE2*, *APOE3*, *APOE4*, sex; male (ochre fill) and female (blue fill), and age; young (alpha 0.2), middle age (alpha 0.6), old (alpha 1). *Statistics:* data from individual experiments were Z-scored. For the “Aging” plot, linear mixed-effects model was fitted for each variable using *sex*, *age*, and *APOE* genotype as fixed effect. Wald χ² tests (HC3-robust) evaluated three-way *sex*×*APOE*×*age* interaction effects. For the individual plots within age group young, middle age and old, linear mixed-effects model was fitted with *sex* and *APOE* genotype as fixed effects. Wald χ² tests (HC3-robust) evaluated the interaction effect of *sex*×*APOE*, male within *APOE* comparisons, female within *APOE* comparisons, *sex* differences within each *APOE* genotype. P-values were FDR-corrected across contrasts, p<0.05: *, p<0.01:**, p<0.001:***.. N = 5 animals per sex, age, *APOE* genotype.

LY particles in the hippocampus were also not affected by a three-way interaction of *age* x *sex* x *APOE* (**Fig. 10C-D**). Pairwise comparisons did not show differences in total LY levels, but N-LY levels in young *APOE4* females were increased compared to *APOE2* – equivalent to the cortex (**Fig. 9C**). Total-LY-Ch levels were specifically regulated at old age; while *APOE4* males showed decreased levels compared to *APOE2* males, *APOE4* females had increased T-LY-Ch levels comparatively (**Fig. 9D**). This effect was also observed on N-LY-Ch levels, but with only an increase in *APOE4* females compared to males. This demonstrates that in the hippocampus at old age there is a specific increase in total and neuronal LY-Ch in *APOE4* females.

Lastly, total LAMP1 LY’s showed a three-way interaction effect of *sex* x *age* x *APOE* genotype, dependent on the difference between *APOE2* and *APOE3* in EC (**Fig. 10E**). In females, there was a steady increase of T-LY levels with age in the *APOE2* and *APOE4* genotype, whereas there was a steady decline with age in *APOE3* females in the EC. In *APOE2* and *APOE4* males, there was a decline with age, in contrast to their female counterparts. *APOE3* males showed an increase in LY’s at middle age, followed by a decrease in old age, following the curve of a sharp peak, instead of a steady decline. Neuronal-LY levels were not affected by the three-way interaction; however, a pairwise contrast at the young age group showed a significant increase in N-LY’s in *APOE4* females versus *APOE2* and *APOE3*, while there were no significant differences at middle and old age.

There were no differences in total LY-Ch levels in the EC (**Fig. 10F**). However, pairwise contrasts showed neuron specific LY-Ch levels at old age increased in *APOE2* males and females compared to *APOE3* males and *APOE4* females respectively. Notably, in every genotype at old age, there was a female-specific increase in N-LY-Ch, compared to males. Interestingly, starting at middle age N-LY-Ch levels in males and females started to diverge, which was significant at old age, in the EC. N-LY-Ch levels were the highest in *APOE2* females (followed by *APOE2* males), raising the question of whether increased N-LY-Ch at old age may be a protective mechanism in relation to AD risk.

## DISCUSSION

In this study, we performed a comprehensive analysis of intracellular cholesterol trafficking in the mouse brain, with a specific focus on the impact of *APOE* genotype, sex, and aging on intracellular cholesterol, endosomal and lysosomal compartments (EE, LE, LY), and their cholesterol content (EE-Ch, LE-Ch, LY-Ch). High magnification confocal microscope images of three different brain regions were included: cortex (CTX), hippocampus (HIPP), and entorhinal cortex (EC). Using robust statistical analyses, we assessed the main effects of the variables *APOE* genotype, *sex* and *age*, and by grouping or segregating data, we were able to assess interaction effects, or simple effects within a certain group.

### Absence of effects on total cholesterol

One of the primary goals of this study was to assess the effect of *APOE* genotype, *sex* and *age* on D4H*-labeled cholesterol spots. The only significant main effect on total cholesterol levels was exerted by sex; female mice had significantly more total and neuronal cholesterol in the cortex compared to male mice. Although it has been shown that female mice rely more heavily on estrogen signaling in the hippocampus for certain cognitive tasks (Rinaudo et al., 2022), there are no studies specifically looking into cholesterol levels. The difference we observed, however, was specific to the cortex, even though the other brain regions may be suggestive of a similar trend to increased cholesterol in females. This may be in line with an MRI study in human participants that showed female participants had increased cortical thickness (Ritchie et al., 2018).

### APOE genotype effects at young age

At young age, EE, LE, and LY levels were often similar across brain regions, whereas these patterns diverged with aging. Neuronal LE levels decreased with *APOE4* (*APOE2* > *E3* > *E4*) in the young mice, while neuronal LY levels showed the opposite pattern (*APOE2* < *E4*). This effect was not significantly identified in each brain region; however, plots from all three brain regions show a similar pattern. This may indicate that at young age, there is a decrease in cholesterol uptake, but that there is a cholesterol trafficking impairment in lysosomes, leading to increased lysosomal cholesterol. This effect virtually disappeared at middle and old age, though this may be due to diverging patterns in the male and the female mice with aging.

The largest effect was observed in the young EC, with neuronal LE’s, EE’s, as well as total/neuronal EE-Ch and LE-Ch decreasing in *APOE4<APOE2* (while *APOE3* levels were in between). Neuronal LY levels trended to increase with *APOE4*, but total and neuronal LY-Ch levels were not affected at young age. These effects leveled out with age, except specificly in *APOE4* mice, where neuronal LY-Ch decreased. This indicates that neurons in the EC may compensate for the decreased neuronal EE and LE cholesterol metabolism in *APOE4* cells at young age, leading to a decline in LY functionality in cholesterol metabolism at older age.

### The female sex is the strongest determinant of neuronal lysosomal cholesterol levels

The only main effect of sex on any of the endolysosomal or endolysosomal cholesterol metrics was measured in total and neuronal LY-Ch in the EC, driven by increased levels in females. This effect was strongest in neuronal LY-Ch specifically, indicating that the female sex significantly alters lysosomal cholesterol processing in the EC. Broken down by age, this effect was specifically evident at old age in the EC, where females consistently had higher levels of N-LY-Ch compared to males, across *APOE* genotypes. LY levels also followed opposing trajectories in males and females with age; whereas LY’s in females steadily rose with age, they steadily declined with age in males. At middle-age, male and female N-LY-Ch levels started to diverge (with more variability in the *APOE4* genotype), which was in line with the increase in T-LY-s in females with age in both *APOE2* and *APOE4* genotype (opposite to the males). This indicates that aging and the female sex are the strongest determinants of neuronal LY-Ch levels specifically. The divergence with age coincides with estrogen decrease in female mice, occurring around 12-15 months of age, roughly parallel to human perimenopause (Balough et al., 2024). These findings suggest that it is imperative for research into pathological mechanisms relating to lysosomal cholesterol dysfunction in neurodegenerative diseases consider sex as biological variable.

### Young age drives early endosomal function conservation in APOE4 males, but not in females

We observed a highly specific *APOE* genotype effect on neuronal EE-Ch levels in the EC, driven by a decrease in *APOE3* females. Both *APOE4* males and females, as well as *APOE3* females expressed similarly low N-EE-Ch levels, whereas *APOE2* males and females, as wells as *APOE3* males, expressed higher levels, in the EC only. Similar but more significant expression levels were observed in young mice, both in the EC and hippocampus. Therefore, the *APOE* x *sex* effect is driven by young age. Notably, there was a significant decrease in total and neuronal EE and EE-Ch levels in *APOE4* males, compared to *APOE3* and *APOE4*, with effectively similar levels in the EC and hippocampus. Eventually in older age, N-EE-Ch levels in older *APOE4* females increased, and *APOE3* females did not. Interestingly, the *APOE4* female total and N-EE-Ch levels increase was observed across all brain regions. This indicates that effects at young age did not specifically drive the female *APOE4* specific N-EE-Ch increase in older females, and that this old age phenotype was present across brain regions. On the contrary, at middle age, there was no difference, and at old age, total and neuronal EE and EE-Ch levels remained low in *APOE4* males, specifically in the EC and hippocampus. This may indicate that in males, there is a specific early endosome-mediated protective mechanism, starting at young age, that conserves EE and EE-Ch functions, maintaining its utility in older age. It is well known that endosomal uptake of toxic peptide species is part of the development of β-amyloid accumulation in neurons. Therefore, mediating endosomal uptake level throughout aging may be protective. The striking difference here is also the specificity of the effect to the EC and hippocampus regions, possibly indicating that regulating early endosomal functions in males are more likely to be associated with cognitive regulation.

### CD63 late endosomes effects

The effect of age x sex x *APOE* genotype was less well defined in late endosomes and LE-Ch. Specifically, in the EC there were interaction effects on total and neuronal LE-Ch levels, indicating that these levels fluctuate with *APOE* genotype, based on *sex*, with *aging*. In the cortex, there were neuron-specific decreases in LE’s and LE-Ch in *APOE4* females (similar to N-EE and N-EE-Ch levels in the EC), while these effects dissolved with age. In the EC, a similar decrease in N-LE-Ch in young age was observed, and interestingly, at middle age there was a difference in total LE-Ch levels, where levels were highest in *APOE4* females, and steadily decreased in male and female *APOE3* and *APOE4*. However, T-LE-Ch levels were highly decreased in *APOE2* males. This effect was specific to the EC, but again leveled out to no significant differences at old age. It is possible that there were less defined patterns in LE and LE-Ch levels because these organelles may overcompensate for EE functions, or LY functions, adapting to its fellow organelles rather than being directly affected by the variables of *age* x *sex* x *APOE* genotype.

## Key Takeaways

- At young age, the imbalance of endolysosomal compartments and cholesterol, regulated by *APOE4*, likely indicates an impairment in cholesterol metabolism.
- *APOE* genotype predominantly exerts its effects in the EC and hippocampus.
- The female sex is the strongest determinant of LY-Ch, but more so N-LY-Ch levels in the EC, while the interaction of female sex with old age is the strongest determinant of LY levels.
- N-EE and N-EE-Ch levels remain low throughout life in *APOE4* males, while these increase with age in *APOE4* females.
- The entorhinal cortex emerges as a region particularly sensitive to age × sex × *APOE* effects on endolysosomal cholesterol trafficking.

## ACKNOWLEDGEMENTS

This work was supported by a grant from the NIA to T.N. (R01 AG070202).

## Notes

### Competing Interest Statement

The authors have declared no competing interest.

