## Supplemental Information for "Investigating the Effects of *APOE* Genotype on Intracellular Cholesterol and the Endolysosomal System in the Aging Mouse Brain"

### FIGURE LEGENDS

**Supplemental Figure 1. Cholesterol levels by *APOE* genotype and *age*. A)** Total levels of cholesterol represented by box plots as total number of D4H* spots normalized to DAPI nuclei. Comparisons per brain regions, cortex, hippocampus, EC. Genotypes *APOE2*, *APOE3*, *APOE4*. **B)** Neuronal levels of cholesterol box plots representing neuronal D4H* spots normalized for the NeuN neuronal volume. **C)** Total levels of cholesterol represented by box plots as total number of D4H* spots normalized to DAPI nuclei. Age groups: Young, Middle age, Old age. **D)** Neuronal levels of cholesterol box plots representing neuronal D4H* spots normalized for the NeuN neuronal volume. Statistics: data from individual experiments were Z-scored before pooling. Linear mixed-effects model was fitted for each variable using *sex*, *age*, and *APOE* genotype as fixed effects. Estimated marginal means for *APOE* (**A-B**) or *age* (**C-D**), and pairwise comparison between *APOE* or *age* respectively. No significant main effect, not pairwise comparison difference was observed.

**Supplemental Figure 5. The interaction effect of sex x age on early endosomes (EE), late endosomes (LE), lysosomes (LY), and cholesterol associated with EE's, LE's, and LY's.** Data in **A)** and **B)** are from the cortex, data in **C)** and **D)** are from the hippocampus. **A)** and **C)** Box plots representing total number of Rab5 spots, CD63 spots, and LAMP1 particles normalized to number of DAPI nuclei. *Lower row*: box plots representing neuronal number of Rab5 spots, CD63 spots, and LAMP1 particles normalized to the total NeuN neuronal volume. The same metrics but different markers are depicted in plots **B)** and **D)** Box plots representing total Rab5, CD63 and LAMP1 colocalized D4H* spots, normalized to the total number of D4H* spots and subsequently normalized to the number of DAPI nuclei. *Lower row*: box plots representing neuronal number of Rab5, CD63, and LAMP1 colocalized D4H* spots, to normalized to the total number of D4H* spots, and subsequently to the total NeuN neuronal volume. Data was grouped by age; young, middle age, old, and sex; male (ochre fill) and female (blue fill). *Statistics:* data from individual experiments were Z-scored. Linear mixed-effects model was fitted for each variable using *sex*, *age*, and *APOE* genotype as fixed effects, and the interaction effect between *sex* and *age*. Estimated marginal means were then computed for the *age* × *Sex* interaction. Significant *age* x sex interaction effect depicted in grey overlay curve (female), and black overlay curve (male). Significance bars in grey show *age* genotype pairwise comparisons, and in black show *sex* dependent comparisons, $p<0.01$:**. N = 5 animals per sex, age, *APOE* genotype.

Supplemental Figure 1A-B. Cholesterol levels by *APOE* genotype

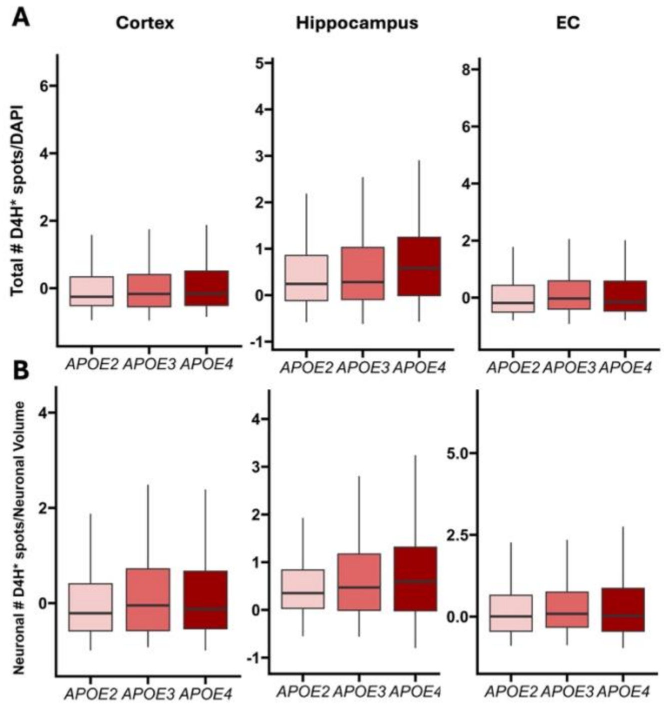


Supplemental Figure 1C-D. Cholesterol levels by age

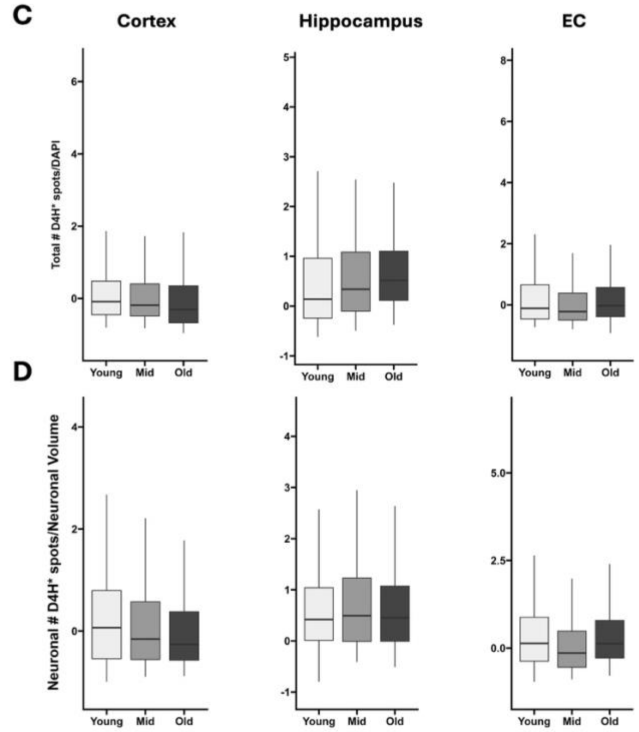

**Supplemental Figure 2A,D. The main effect of *APOE* genotype on early endosomes (EE) and cholesterol associated with EE's**

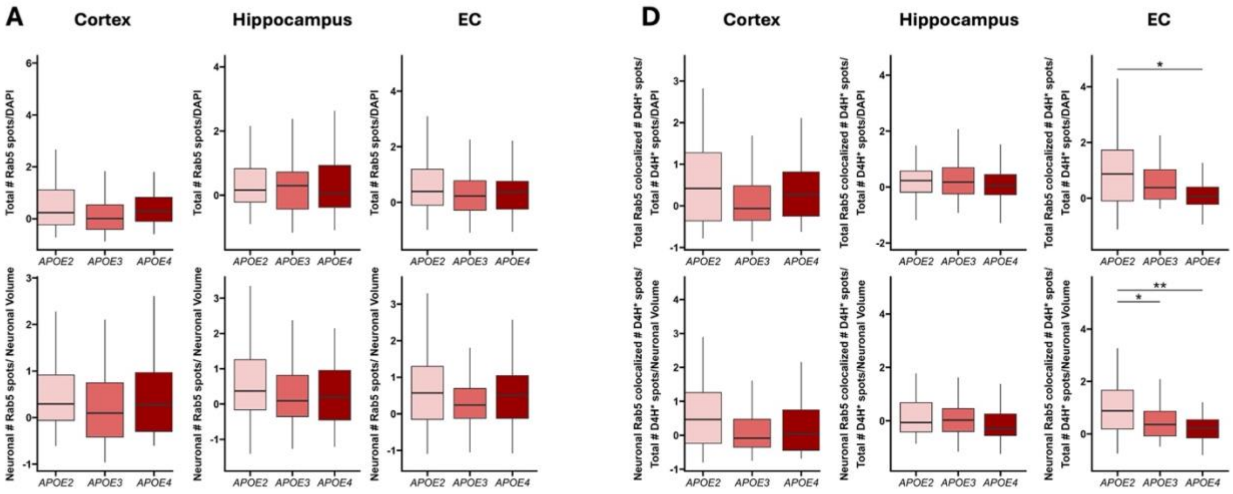


**Supplemental Figure 2B,E. The main effect of *APOE* genotype on late endosomes (LE) and cholesterol associated with LE's**

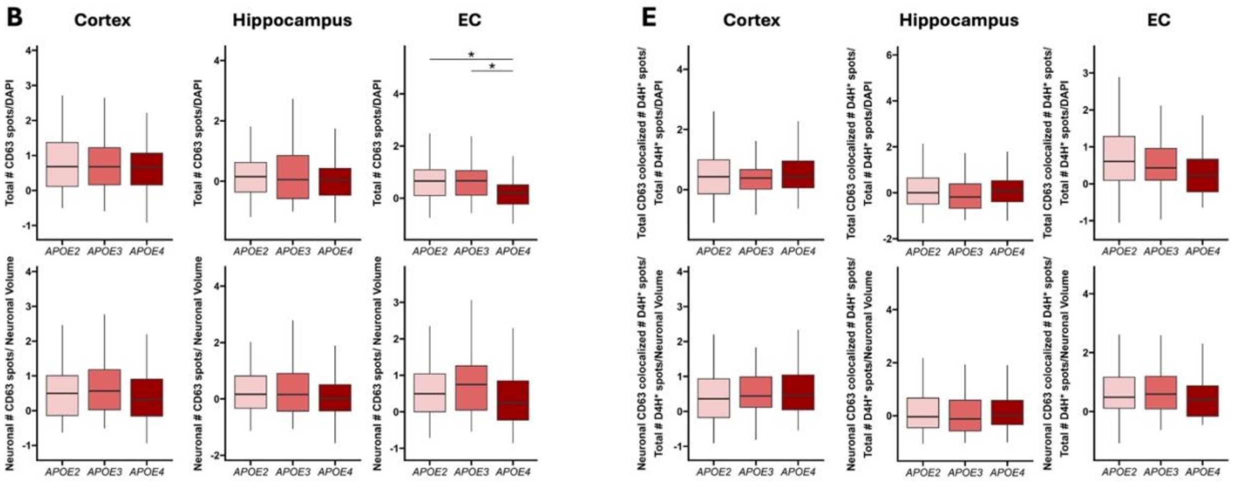

**Supplemental Figure 2C,F. The main effect of *APOE* genotype on lysosomes (LY) and cholesterol associated with LY's**

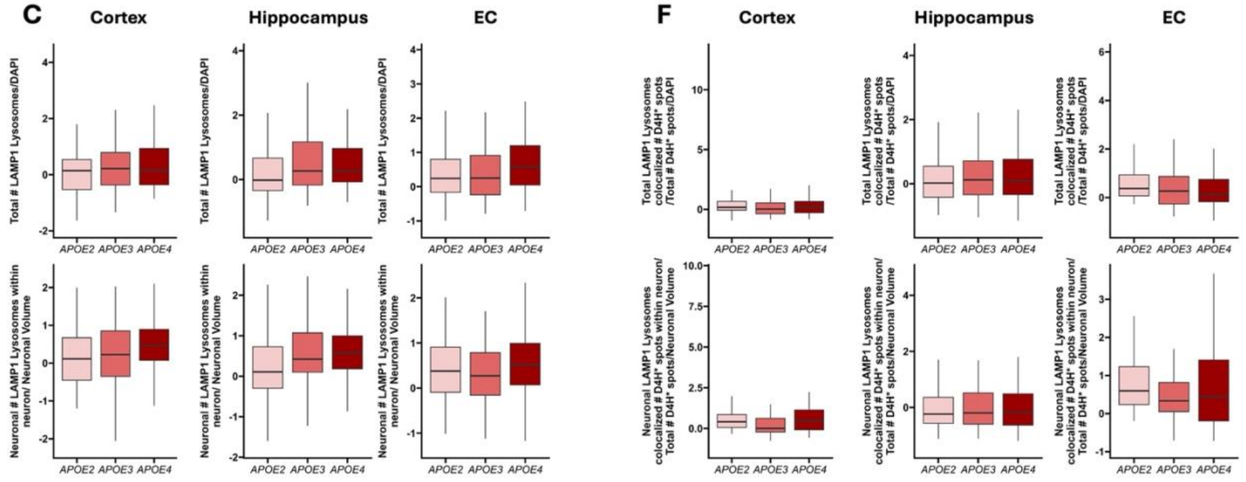

**Supplemental Figure 3A,F. The effect of *APOE* genotype in middle aged mice, in each brain region on early endosomes (EE), late endosomes (LE), lysosomes (LY), and cholesterol associated with EE's, LE's, and LY's in the <u>Cortex</u>**

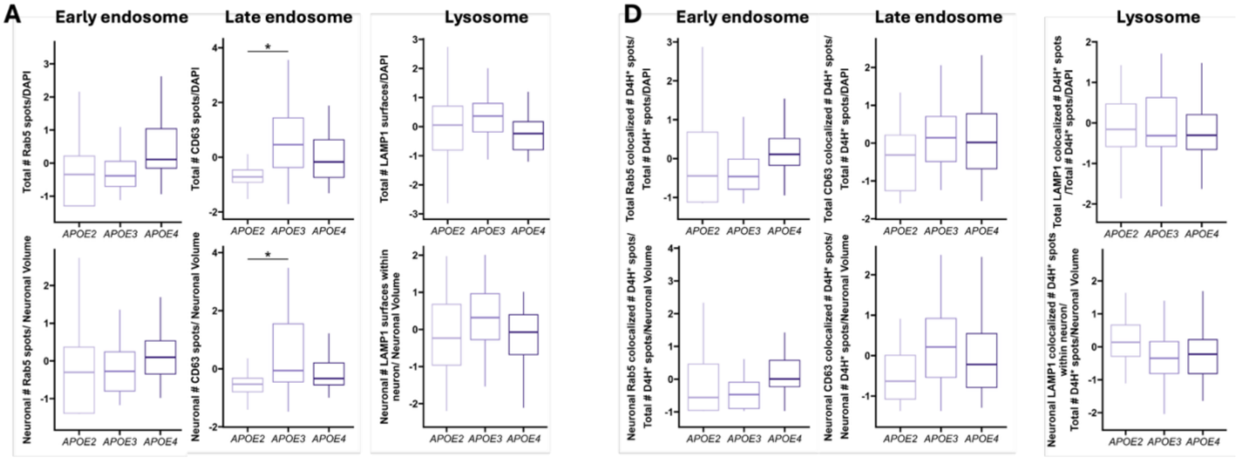


**Supplemental Figure 3B,E. The effect of *APOE* genotype in middle aged mice, in each brain region on early endosomes (EE), late endosomes (LE), lysosomes (LY), and cholesterol associated with EE's, LE's, and LY's in the <u>Hippocampus</u>**

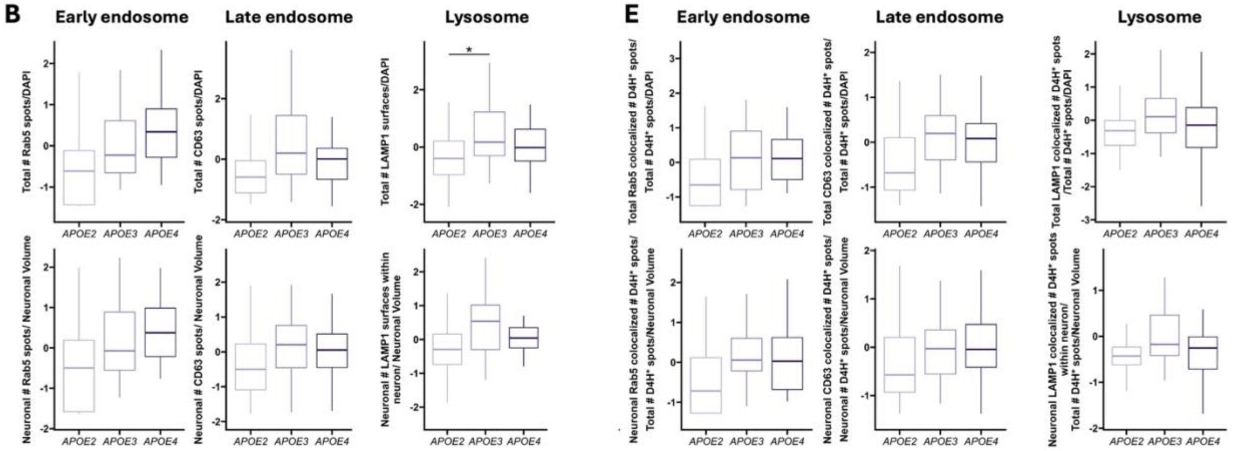

**Supplemental Figure 3C,F. The effect of *APOE* genotype in middle aged mice, in each brain region on early endosomes (EE), late endosomes (LE), lysosomes (LY), and cholesterol associated with EE's, LE's, and LY's in the <u>EC</u>**

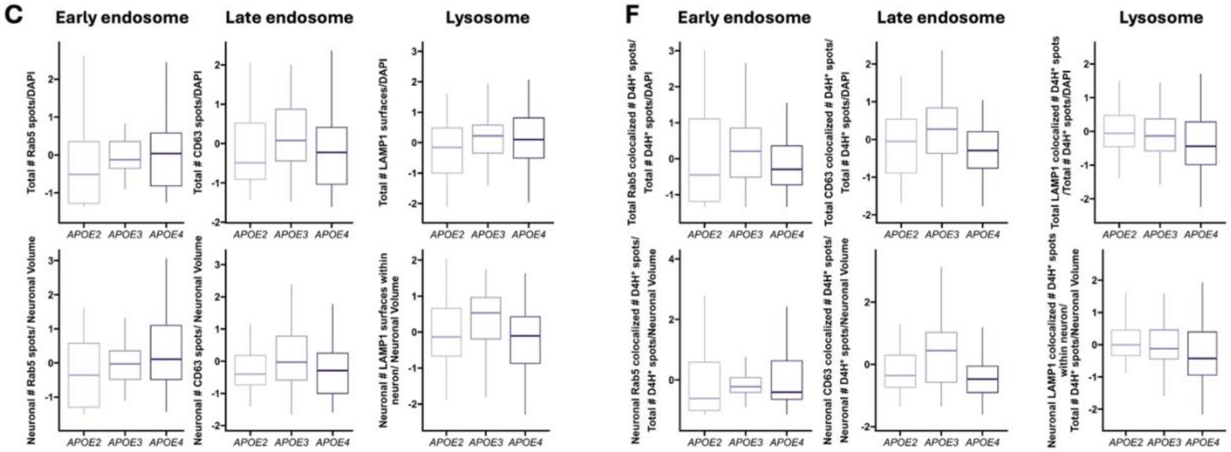

**Supplemental Figure 4A,D. The effect of *APOE* genotype in old mice, in each brain region on early endosomes (EE), late endosomes (LE), lysosomes (LY), and cholesterol associated with EE's, LE's, and LY's in the <u>Cortex</u>**

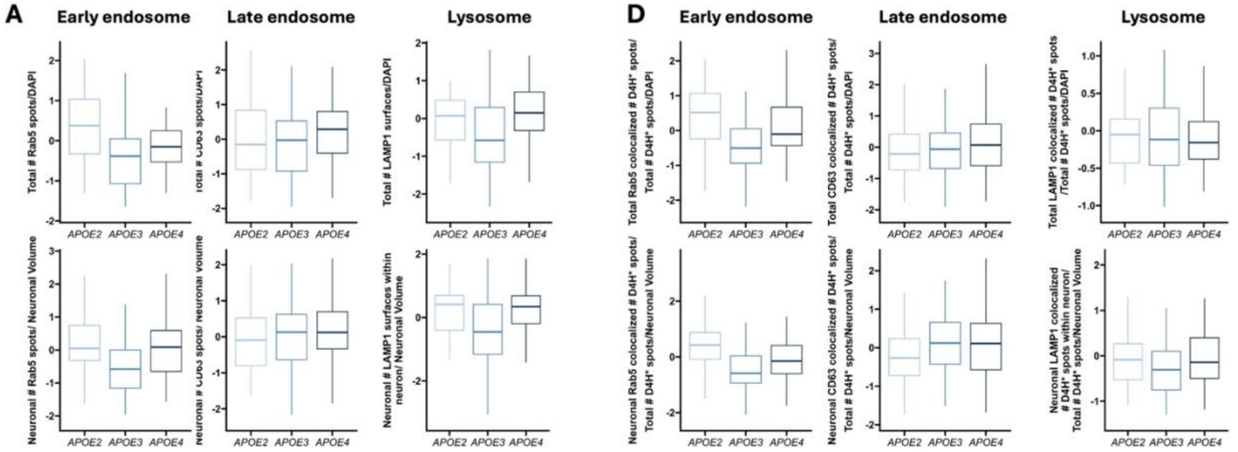


**Supplemental Figure 4B,E. The effect of *APOE* genotype in old mice, in each brain region on early endosomes (EE), late endosomes (LE), lysosomes (LY), and cholesterol associated with EE's, LE's, and LY's in the <u>Hippocampus</u>**

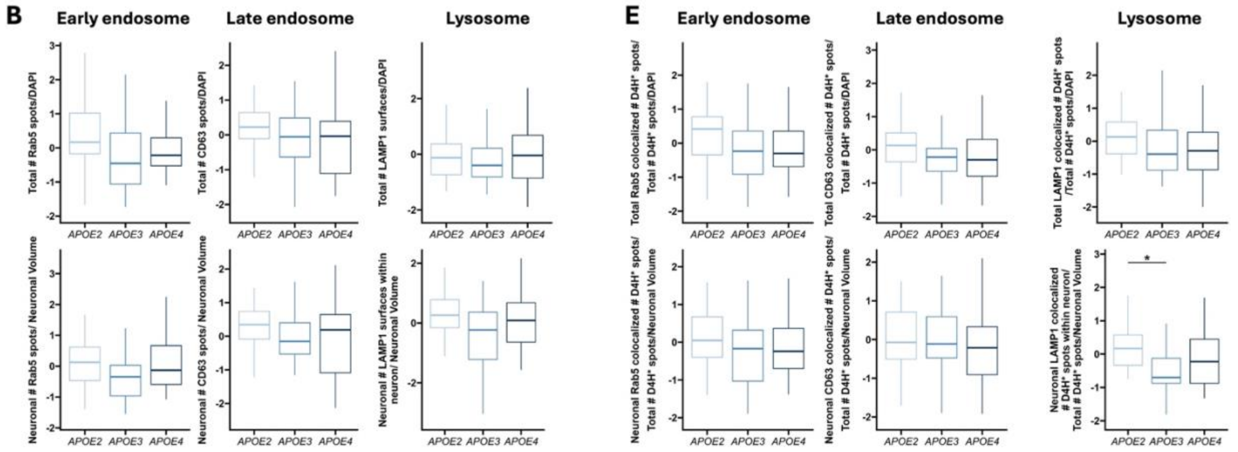

**Supplemental Figure 4C,F. The effect of *APOE* genotype in old mice, in each brain region on early endosomes (EE), late endosomes (LE), lysosomes (LY), and cholesterol associated with EE's, LE's, and LY's in the EC**

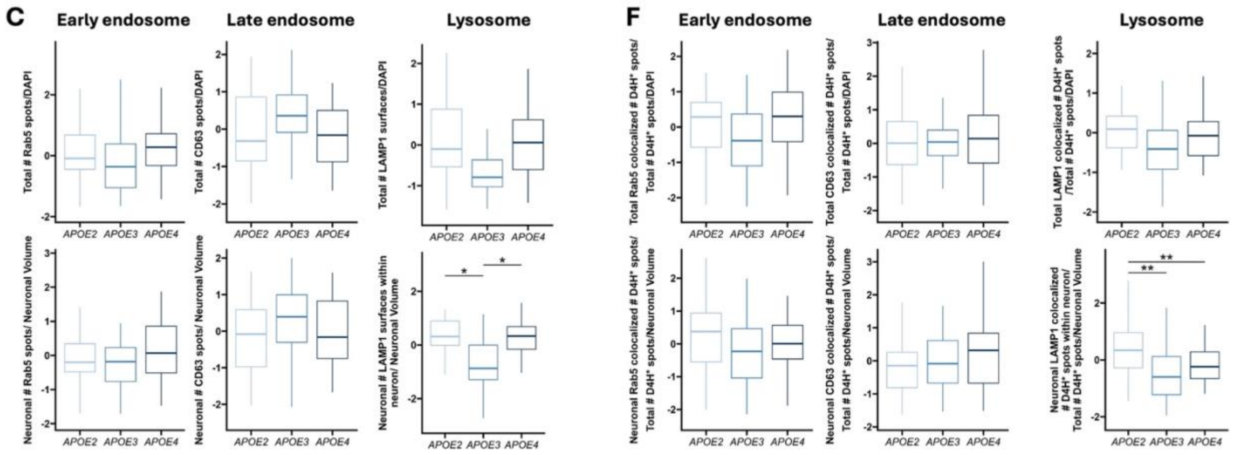

**Supplemental Figure 5A,B. The interaction effect of sex x age on early endosomes (EE), late endosomes (LE), lysosomes (LY), and cholesterol associated with EE's, LE's, and LY's in the <u>Cortex</u>**

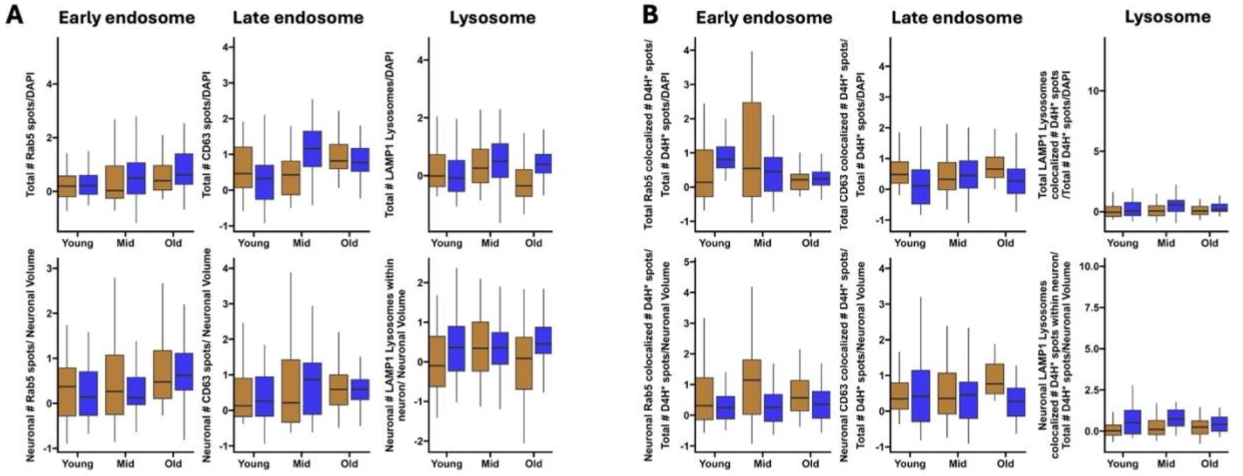


**Supplemental Figure 5C,D. The interaction effect of sex x age on early endosomes (EE), late endosomes (LE), lysosomes (LY), and cholesterol associated with EE's, LE's, and LY's in the <u>Hippocampus</u>**

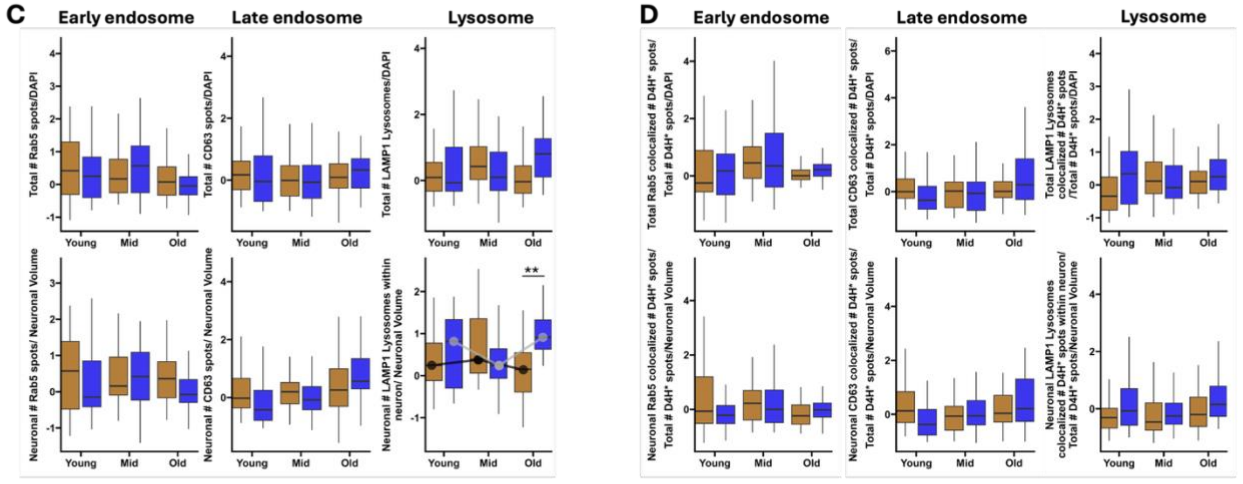
